# Identification of transcription factors involved in the specification of photoreceptor subtypes

**DOI:** 10.1101/2021.11.26.470161

**Authors:** Juan Angueyra, Vincent P. Kunze, Laura K. Patak, Hailey Kim, Katie S. Kindt, Wei Li

## Abstract

During development, retinal progenitors navigate a complex landscape of fate decisions that results in the generation of unique cell types necessary for proper vision. Here, we aim to provide the resources and techniques required to identify factors that are critical for fate decisions in photoreceptors. These factors help create a diversity of photoreceptor subtypes that sustain vision in day and night, enable the discrimination of colors, facilitate the detection of prey and predators, and support other aspects of vision. First, we generate a key resource: a high-quality and deep transcriptomic profile of each photoreceptor subtype in zebrafish. We make this resource openly accessible, easy to explore and integrate it with other currently available photoreceptor transcriptomic datasets. Second, using our transcriptomic profiles, we derive an in-depth map of expression of transcription factors in photoreceptors—potential key players in cell-fate decisions. Third, we explore CRISPR-FØ screening as a fast, efficient and versatile technique to assess the involvement of candidate transcription factors in photoreceptor subtype-specification. We first show that known phenotypes can be easily replicated: loss of S cones in *foxq2* mutants and loss of rods in *nr2e3* mutants. We then explore four additional transcription factors of unknown function (Skor1a, Sall1a, Lrrfip1a and Xbp1) and find no evidence for their involvement in photoreceptor-subtype specification. Finally, we identify novel functions of Tbx2, demonstrating that it plays a central role in controlling the identity of all photoreceptor sub-types within the retina. Our study provides an open roadmap to discover additional factors involved in this process. This dataset and screening method will be a valuable way to explore the genes involved in many essential aspects of photoreceptor biology.

## Introduction

The specification of cell types is a fundamental and complex process that underlies the formation of tissues and organs. Cell specification is especially critical for the primary sensors in sensory systems. Primary sensory cells not only have to acquire distinct and elaborate specializations to detect and transduce physical stimuli, but also must wire with specific circuits to faithfully relay sensory information. In the retina, for instance, photoreceptor specification must coordinate with the distinct developmental timelines of other retinal cell types to form functional visual circuits (1). In addition, photoreceptors have clearly defined subtypes which differ in spectral sensitivity, morphology, density across the retina, and wiring. Comprehensively, the factors required to specify photoreceptor subtypes remains unclear. Our aim is to identify these factors by exploiting technical advantages of the zebrafish model.

The correct specification of photoreceptor subtypes is critical for proper vision. Foundational work across vertebrates has identified transcription factors—*e*.*g*. CRX, OTX2/OTX5, and NEUROD1—that are required to generate photoreceptor progenitors during development, before subtypes are clearly defined (2–7). Numerous studies have investigated how photoreceptor subtypes are specified from these progenitors and have shown that, in mice, the transcription factor NRL and its downstream effector NR2E3 are required for rod specification (8–10). In humans, mutations in NR2E3 cause *enhanced S-cone syndrome*, where failures in rod specification ultimately lead to impaired visual acuity, abnormal color vision, night blindness and retinal degeneration (11). In addition to conserved factors involved in rod specification, THRB is required for L-cone specification in mice (12), zebrafish (13) and, most likely, birds (14). In the absence of these “key gatekeepers”—NRL, NR2E3 or THRB—mouse photoreceptor progenitors acquire an S-cone fate, and S-cone specification is commonly considered a passive process (7). Yet, mounting evidence—derived mainly from work species other than mouse—challenges this simplistic model. For example, while Nr2e3 is also required to generate rods in other vertebrates (15), Nrl is dispensable (16–18). In addition, UV- and S-cone specification in zebrafish is far from passive and requires the action of Tbx2b and Foxq2, respectively (19, 20). Together these studies highlight that our understanding of how photoreceptor subtypes are specified remains incomplete.

Our study seeks to provide the resources and methods required to efficiently identify additional genes involved in photoreceptor specification. The study is divided into four sections. First, we obtain deep and high-quality transcriptomic profiles (RNA-seq) of the five zebrafish photoreceptor subtypes. Second, we explore this RNA-seq dataset and identify multiple transcription factors that could potentially be involved in the specification of photoreceptor subtypes. Third, we show that a CRISPR-based FØ-screening approach (21, 22) is a reliable platform to identify transcription factors involved in photoreceptor specification. We benchmark our screening method by replicating known phenotypes of *foxq2* and *nr2e3* mutants: FØ larvae that carry mutations in *foxq2* lose S cones while those that carry mutations in *nr2e3* lose rods (15, 20). Subsequently, we explore the role of four additional transcription factors with no known function (Skor1a, Sall1a, Lrrfip1a, Xbp1) and find that they are not critical to specify photoreceptor subtypes. Finally, we demonstrate the potential of this platform by describing novel roles for the transcription factors Tbx2a and Tbx2b in photoreceptor specification, demonstrating that UV-cone specification requires both Tbx2a and Tbx2b, and that Tbx2a and Tbx2b, respectively, maintain the identity of L cones and S cones by repressing M-cone cell fate.

Our dataset and methods can be applied to further our understanding of how photoreceptors acquire their final identities and to explore other important aspects of photoreceptor biology that also differ between subtypes (phototransduction, metabolism, synaptic wiring, *etc*.). To facilitate such future studies, we provide open and easy access to our transcriptomic dataset and analysis, and integrate it with other relevant and available datasets (https://github.com/angueyraNIH/drRNAseq/). Ultimately, the knowledge gained by exploring these datasets can be used to inform strategies to control the photoreceptor differentiation in organoids—a potential gateway for cell-replacement therapies in retinal degenerations.

## Results

### Transcriptomic analysis of adult zebrafish photoreceptors

Identifying the transcription factors required to specify photoreceptor subtypes is critical for understanding the normal development of the retina and to inform cell-replacement therapies to restore vision. RNA-seq is a powerful way to identify novel genes expressed in cell subtypes. Although RNA-seq approaches have been used to identify genes differentially expressed between photoreceptor sub-types in many species, the limited transcriptome depth derived from single-cell techniques (23) constitutes a barrier in the reliable detection of transcription factors, which are frequently expressed at low levels (24). To obtain a deep, highquality RNA-seq dataset from zebrafish photoreceptors, we used well-characterized transgenic lines that express fluorescent proteins in each subtype with high specificity, including rods—*Tg(xOPS:GFP)*, UV cones—*Tg(opn1sw1:GFP)*, S cones—*Tg(opn1sw2:GFP)*, M cones—*Tg(opn1mw2:GFP*) and L cones—*Tg(thrb:tdTomato*) (Figure 1A) (13, 25–28), to manually collect pools of photoreceptors of a single subtype under epifluorescence (29). Manual collection allowed us to focus on fluorescent and healthy photoreceptors, with intact outer segments, cell bodies and mitochondrial bundles, and to avoid cellular debris and other contaminants (Figure 1B). For each sample, we collected pools of 20 photoreceptors of a single subtype derived from a single adult retina. After collection, we isolated mRNA and generated cDNA libraries for sequencing using SMART-seq2 technology (Figure 1C). In total, we acquired 6 rod samples and 5 UV-, 6 S-, 7 M-and 6 L-cone samples. On average, we were able to map 86.4% of reads to the zebrafish genome (*GRCz11*; range: 76.3% - 90.4%), corresponding to 10.19 million ± 1.77 million mapped reads per sample (mean ± s.d.) and to an average of 7936 unique genes per sample (range: 5508 – 10,420) (Figure 1D). This high quantity of reads and unique genes demonstrates that our technique provides substantially deep transcriptomes—especially when compared to single-cell droplet-based techniques where the number of reads per cell is largely less than 20,000, corresponding to more than 2000-fold differences in depth (23, 30, 31). Using unsupervised clustering (t-distributed Stochastic Neighbor Embedding or tSNE), we found that samples correctly clustered by the subtype they were derived from. Proper clustering provides evidence that differences in gene expression captured in our RNA-seq data stem mainly from distinctions between photoreceptor subtypes (Figure 1E).

**Figure. 1.**
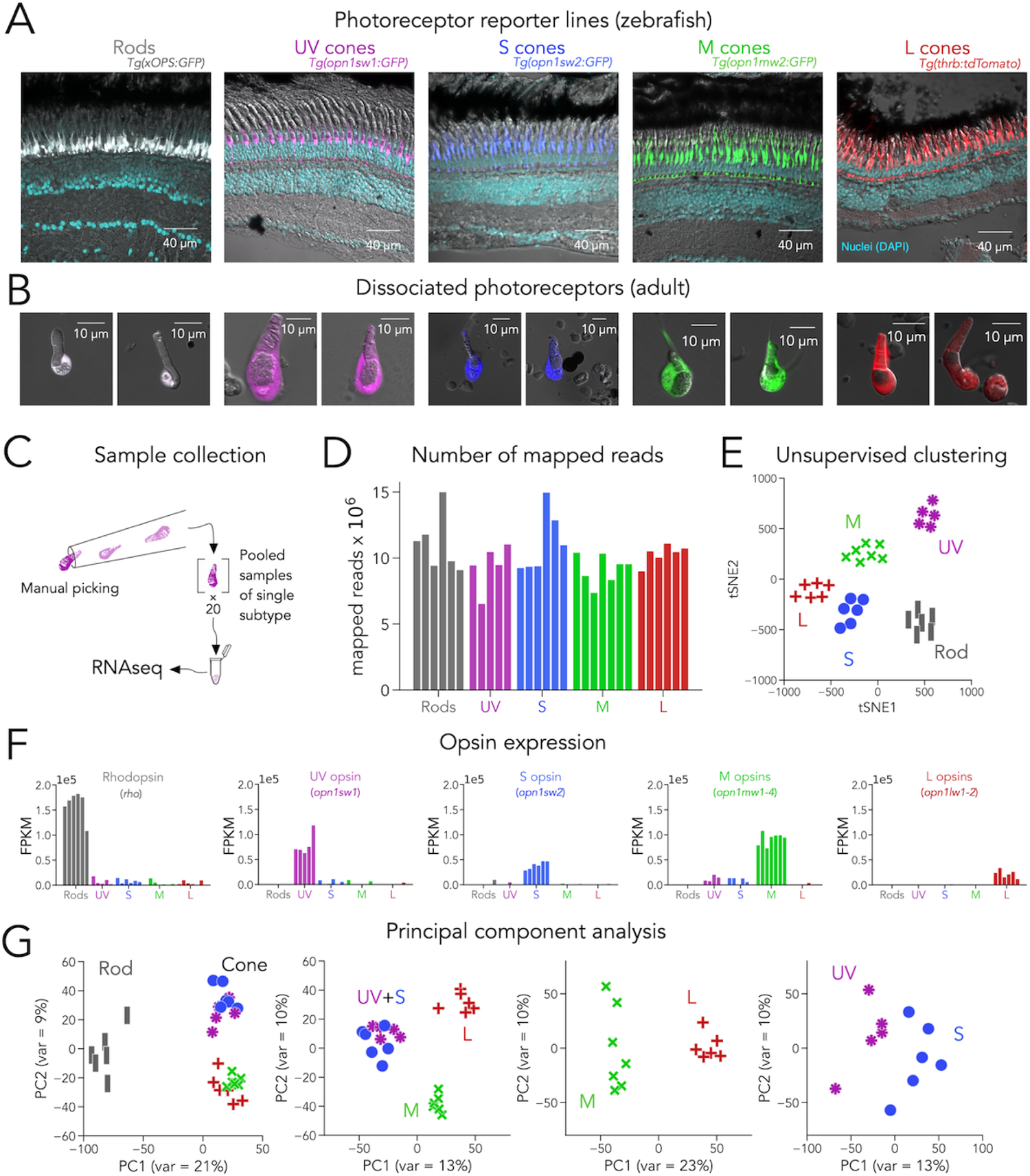
Transcriptomic profiling (RNAseq) of zebrafish photoreceptors. **(A)** Confocal images of fixed adult zebrafish retinal cross-sections, from transgenic reporters used to identify photoreceptor subtypes. Reporter expression is exclusive to the outer retina, and each line labels a single photoreceptor subtype with unique morphology, including rods (grey), UV cones (magenta), S cones (blue), M cones (green) and L cones (red). The inner retinal layers can be distinguished in the overlayed nuclear stain (DAPI, cyan) and transmitted DIC image (grey). **(B)** Confocal images of dissociated and live photoreceptors of each subtype, identified by fluorescent reporter expression. Photoreceptors have preserved outer segments and identifiable mitochondrial bundles. **(C)** Sample collection method. After dissociation, 20 healthy photoreceptors of a single subtype were identified by fluorescence and manually picked with a glass micropipette and pooled as a single RNAseq sample. **(D)** High transcriptome depth shown by the number of reads successfully mapped to the zebrafish genome (GRCz11) for each RNAseq sample. **(E)** Clustering using t-distributed stochastic neighbor embedding correctly separates samples by their original subtype. **(F)** Plots of opsin expression show high counts for the appropriate opsin in each sample (in fragments per kilobase per million reds or FPKM) and low-to-negligible counts of other opsins. For M-opsin quantification, we calculated the sum of counts for *opn1mw1, opn1mw2, opn1mw3* and *opn1mw4* and for L-opsin quantification, we summed counts for *opn1lw1* and *opn1lw2*. **(G)** Iterative principal component analysis (PCA) shows that differences in gene expression separate rods and cones (first panel), and UV/S cones from M/L cones (second panel). M and L cones can also be distinguished by a single principal component (third panel), while separation of UV and S cones is more difficult (fourth panel).

The expression of opsin genes is unique between photoreceptor subtypes and, under normal conditions, a reliable marker of subtype identity. Consistent with this idea, each sample had a high number of reads for the appropriate opsin. Furthermore, there were very low reads for other opsins, corroborating the purity of our samples (Figure 1F). We also found that reads for phototransduction genes were high and consistent with the known differences in gene expression between rods and cones (*e*.*g*., *gnat1*, rod transducin, had high reads only in rod samples while *gnat2*, cone transducin, had high reads in all cone samples) and between cone subtypes (*e*.*g*., expression of *arr3a* in M and L cones and *arr3b* in UV and S cones) (Figure 1 – Figure supplement 1) (31, 32). To expand our analysis to other genes, we first used principal component analysis (PCA) as an unbiased approach to determine how variability in gene expression defines photoreceptor subtypes. PCA revealed that most of the differences in gene expression were between rods and cones. When cones were considered separately, the biggest differences in gene expression arose from two groupings: UV and S cones vs. M and L cones. Subsequent analysis revealed a clear separation of M and L cones, with UV and S cones showing the least differences (Figure 1G). Guided by this analysis, we performed differential gene-expression analysis by making pairwise comparisons following the directions of the principal components, revealing a diverse set of ∼3000 differentially-expressed genes, many of unknown function in photoreceptors (Figure 1 – Figure Supplement 2 and Supplementary Data 1).

In summary, our manual, cell-type specific, SMART-seq2-based approach yielded high-quality zebrafish photoreceptor transcriptomes, with very low contamination and ∼2000-fold more depth than published single-cell RNA-seq studies in the retina, and thus has a particularly high signal-to-noise ratio for differential gene-expression analysis (23, 30, 31, 33) (Figure 1 – Figure Supplement 3). As exemplified by phototransduction proteins (and transcription factors below), our dataset is in good agreement with current knowledge of photoreceptor-expressed genes; in addition, it uncovered novel and unexplored differences in gene expression between photoreceptor subtypes. This RNA-seq dataset constitutes a useful resource to explore genes that are generally or differentially expressed by photoreceptor subtypes which could be involved in multiple aspects of photoreceptor biology, especially when integrated with other relevant studies (see discussion). Our subsequent analyses center on transcription factors.

### Expression of transcription factors in zebrafish photoreceptors

Transcription factors play a critical role in photoreceptor specification. Therefore, we isolated all RNA-seq reads that could be mapped to transcription factors, and detected significant expression of 803 transcription factors. When ranked by average expression levels across all samples, *neurod1* was revealed as the transcription factor with the highest expression by ∼5-fold. High expression in adult photoreceptors suggests that Neurod1 plays a role in the mature retina and not only during development or regeneration (5, 34, 35). Among the 100 most highly-expressed transcription factors, we identified genes clearly established as being critical during photoreceptor development or specification including *crx, otx5, rx1, rx2, nr2e3, six6b, six7, meis1b, egr1, foxq2* and *thrb* (Figure 2A, blue bars) (2–4, 20, 36–41). Only a limited number of studies explore the function of some of the remaining transcription factors in photoreceptors, despite their high expression on our dataset (Figure 2A, grey bars) (42–50). Furthermore, many of these have not been previously studied (Figure 2A, black bars), suggesting that our current knowledge on the control of genetic programs in photoreceptors remains incomplete.

**Figure. 2.**
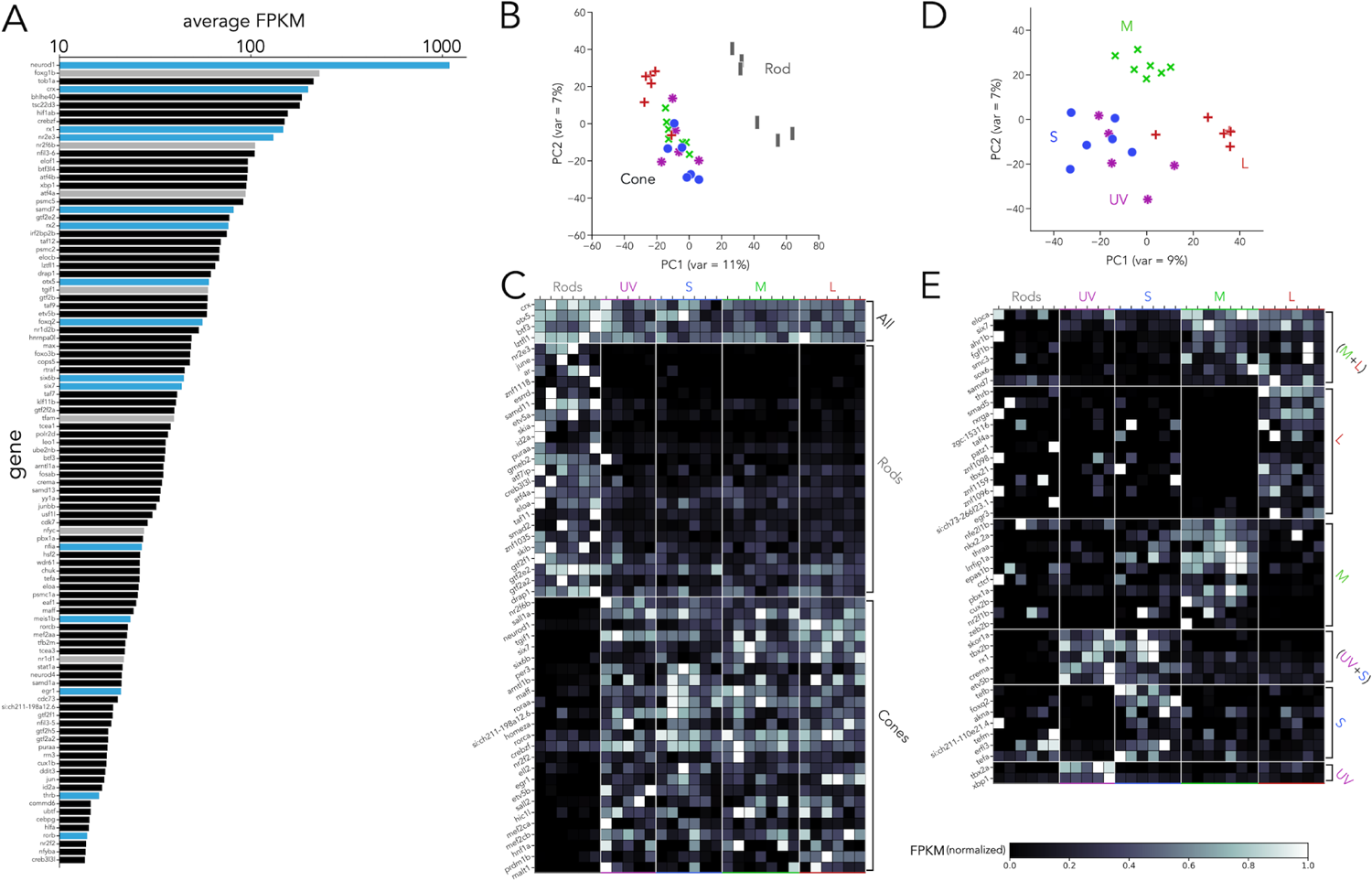
Transcription factor expression in zebrafish adult photoreceptors. **(A)** Top 100 transcription factors ranked by average expression across all samples, displayed on a log-scale. Genes highlighted in blue are known to be critical in photoreceptor progenitors or during photoreceptor development. Some limited information about function in photoreceptors exists for genes highlighted in grey. **(B)** Principal component analysis (PCA) of transcription factors shows that differences between rods and cones are the main source of variance. **(C)** Heatmap showing differential expression of transcription factors between rods and cones and divided into three groups: consistently expressed by all photoreceptors, enriched in rod samples and enriched in cones samples. Grey values indicate expression level normalized in each row by the maximal value. Rod- and cone-enriched genes have been arranged by degree of enrichment. **(D)** PCA of cone samples shows that the largest differences in expression separate L cones (PC1) and M cones (PC2), while separation of UV and S cones is more difficult. **(E)** Heatmap of transcription factors differentially expressed in cone subtypes, divided into 6 relevant groups. Full list of differentially-expressed transcription factors available in Supplementary Data 2 and 3.

Next, we used PCA to understand how the expression of these 803 transcription factors differs between photoreceptor subtypes. Like our whole-transcriptome analysis, we found that most of the differences in transcription-factor expression can be attributed to differences between rods and cones (Figure 2B). By performing pairwise comparisons of transcription factors based on rod vs. cone expression, we identified three relevant groups: (1) consistently expressed across all subtypes, (2) rod-enriched and (3) cone-enriched (Figure 2C). Consistent with previous studies, expression of *crx* and *otx5* was similar across subtypes (2, 3). *nr2e3, samd7* and *samd11* showed clear rod-enrichment (10, 51, 52) and *six6a, six6b, six7, sall1a* and *neurod1* showed cone-enrichment (5, 37, 38, 53). By expanding our analysis beyond previously characterized genes, our dataset revealed a total of 75 transcription factors with significant differential expression between rods and cones, many of which have no clear function in photoreceptors (Figure 2C and Supplementary Data 2).

We next examined the variance in transcription-factor expression between cone subtypes. PCA revealed that both L and M cones could be distinguished by differences in transcription-factor expression alone, while UV and S cones again showed the fewest differences (Figure 2D). By analyzing cone subtypes, we found a total of 47 differentially expressed transcription factors. Seven transcription factors were significantly enriched in both L and M cones compared to UV and S cones (Figure 2E) and included *ahr1b*—a gene associated with Retinitis Pigmentosa in humans (54)—and *six7*—known to be involved in cone progenitor development and survival (37). Twelve were enriched in L cones, including *thrb*—known to be critical for L-cone identity across vertebrates (12, 13)—and *rxrga*—a regulator of L-opsin expression in mouse (55). Amongst the ten M-cone enriched transcription factors, we identified *thraa*—another thyroid hormone receptor, confirmed to be expressed by photoreceptors (56)— and *lrrfip1a*. A small group of just five genes was enriched in both UV and S cones compared to L and M cones and included *tbx2b* and *skor1a*. Seven transcription factors were enriched in S cones—including *foxq2*—and two were enriched in UV cones—*tbx2a* and *xbp1* (Figure 2E and Supplementary Data 3). To validate our RNAseq, we used a fluorescent *in-situ* hybridization assay and detected expression in photoreceptors of several transcription factors identified through this analysis (Figure 2 – Supplement 1).

In summary, our transcriptomic analysis is in good agreement with our current knowledge of transcription factor expression in photoreceptors. Additionally, it reveals novel patterns of expression between photoreceptor subtypes. Notably, a considerable fraction of these transcription factors has no clear function in photoreceptors in zebrafish or in other species, making them clear targets for follow-up studies aimed at understanding photoreceptor specification and other subtype-specific functions.

### FØ screening as a reliable platform to explore the involvement of transcription factors in photoreceptor identity

Our dataset revealed an extensive collection of transcription factors potentially involved in photoreceptor differentiation. Given the high number of candidates and our limited knowledge on their function, we sought to establish methods for efficient identification of transcription factors involved in subtype specification. Instead of attempting to create full germline mutants for each gene—a process that takes 2– 3 generations and 6-12 months—we focused on a fast and highly efficient CRISPR-Cas9-guided mutagenesis approach that enables FØ screening (21, 22). In these FØ screens, single-cell zebrafish embryos are injected with Cas9 protein and multiple guide RNAs (gRNAs) targeting a gene of interest. Just days after injection, in the FØ generation, injected larvae can be assessed phenotypically. Although these FØ larvae are genetic mosaics (some cells may not carry mutations and mutations are not identical in every cell), each larva can be genotyped to establish a robust link between gene function and phenotype. In addition, FØ larvae can be created in the context of any combination of existing transgenic or mutant lines, avoiding the need to maintain additional crosses. For all the analyses presented below, we calculated photoreceptor densities in the central retina at 5 days post-fertilization, using subtype-specific reporter lines (Table 1). All analyses correspond to FØ larvae that have been genotyped to confirm mutations in the targeted gene (see methods), and guides were tested to ensure a high rate of mutations (Table 3).

**Table 1.**
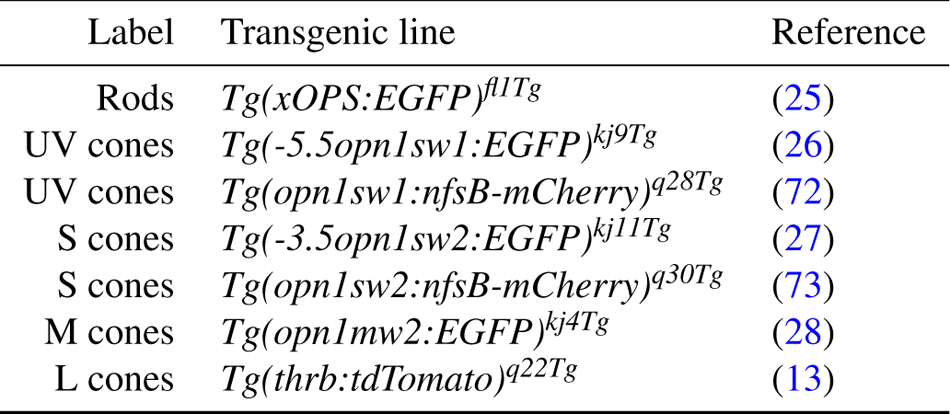
Zebrafish transgenic lines

To benchmark this FØ screen in the context of photoreceptor specification, we first targeted two genes with subtypespecific expression in our RNAseq and that are known to be involved in this process—Foxq2 and Nr2e3. Among transcription factors, *foxq2* is expressed at relatively high levels, ranking 33^rd^ (Figure 2A), and is specifically enriched in S cones, with negligible expression in other photoreceptor subtypes (Figure 3A). Loss-of-function of *foxq2* mutants are characterized by a complete loss of S cones and S-opsin expression, and a slight increase in M-opsin expression (20). For our FØ analysis, we designed two gRNAs targeted against the DNA-binding forkhead domain of Foxq2 (57). Compared to wild-type controls, FØ[*foxq2*] displayed a marked decrease of ∼85% in the density of S cones (Figure 3B and Figure 3– figure supplement 1). Consistent with the slight increase in M-opsin expression in germline loss-of-function mutants, we also found a small but significant increase of ∼24% in the density of M cones in FØ[*foxq2*], while the densities of rods, UV cones, and L cones remained unchanged (Figure 3C).

**Figure. 3.**
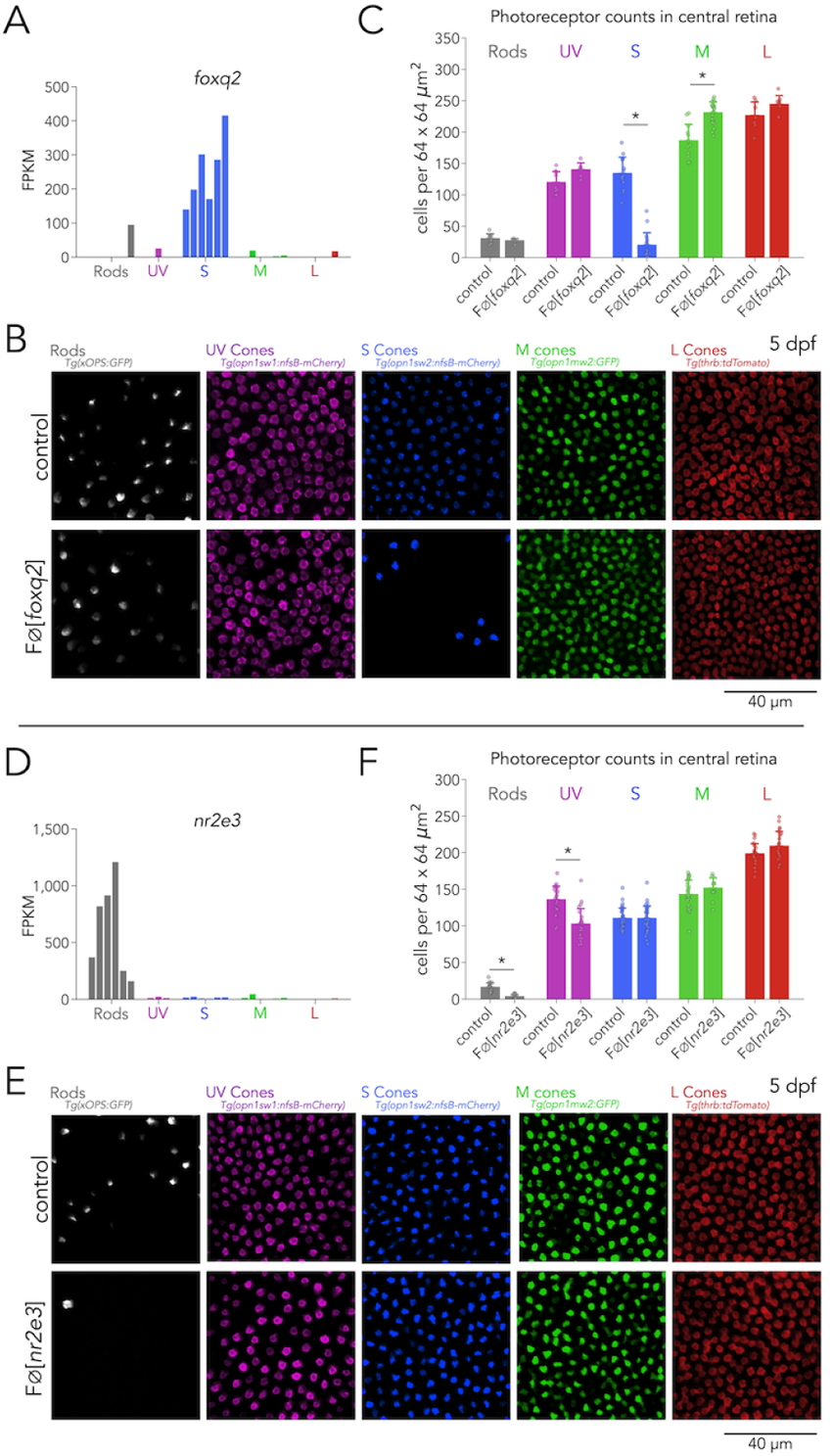
Foxq2 is required for S-cone specification and Nr2e3 for rod specification. **(A)** Expression of *foxq2* shows clear S-cone specificity. **(B)** Mutations in *foxq2* cause a loss of S cones. Representative confocal images of the central retina of control (top row) and FØ[*foxq2*] (bottom row) larvae at 5 dpf. Each column corresponds to a transgenic line that labels a unique photoreceptor subtype, pseudo-colored according to photoreceptor subtype. **(C)** Quantification of photoreceptors in control and FØ[*foxq2*] larvae. Bars represent averages, error bars correspond to standard deviations, and markers correspond to individual retinas. There is a significant ∼85% reduction in S cones in FØ[*foxq2*] compared to *wt* controls (Mann-Whitney U = 221.00, p = 4.12 × 10^−6^, n_wt_ = 13, n_FØ[*foxq2*]_= 17), a smaller but significant ∼24% increase in the density of M cones compared to *wt* controls (Mann-Whitney U = 21.00, p = 9.17 × 10^−5^, n_wt_ = 12, n_FØ[*foxq2*]_= 21) and no significant differences in the densities of rods (Mann-Whitney U = 34.50, p = 0.41, n_wt_ = 9, n_FØ[*foxq2*]_= 6), UV cones (Mann-Whitney U = 11, p = 0.07, n_wt_ = 9, n_FØ[*foxq2*]_= 6), or L cones (Mann-Whitney U = 15, p = 0.15, n_wt_ = 7, n_FØ[*foxq2*]_= 21). **(D)** Expression of *nr2e3* shows enrichment in rods. **(E)** Mutations in *nr2e3* cause a loss of rods. Representative confocal images of the central retina of control (top row) and FØ[*nr2e3*] (bottom row) larvae at 5 dpf. **(F)** Quantification of photoreceptors in control and FØ[*nr2e3*] larvae. Bars represent averages, error bars correspond to standard deviations, and markers correspond to individual retinas. There is a significant ∼80% reduction in rods in FØ[*nr2e3*] compared to controls (Mann-Whitney U = 358.00, p = 2.22 × 10^−7^, n_wt_ = 19, n_FØ[*nr2e3*]_= 19), a smaller but significant ∼25% reduction in UV cones (Mann-Whitney U = 416.00, p = 1.5 × 10^−5^, n_*wt*_ = 24, n_FØ[*nr2e3*]_= 22), and no significant differences in the densities of S cones (Mann-Whitney U = 469.00, p = 0.88, n_*wt*_ = 30, n_FØ[*nr2e3*]_= 32), M cones (Mann-Whitney U = 134.00, p = 0.21, n_*wt*_ = 30, n_FØ[*nr2e3*]_= 12), or L cones (Mann-Whitney U = 193.50, p = 0.12, n_*wt*_ = 24, n_FØ[*nr2e3*]_ = 22).

As a second positive control, we created mutations in *nr2e3*—a rod enriched-gene (Figure 3D), known to be critical for the generation of rods in vertebrates (7, 9, 10)—by injecting two gRNAs targeted against exon 1. As in germline *nr2e3* mutants (15), FØ[*nr2e3*] have a pronounced loss of ∼80% of rods (Figure 3E). Interestingly, in FØ[*nr2e3*] we also identified a ∼25% decrease in UV-cone densities—which has not been previously reported—suggesting an unrecognized role of Nr2e3 in cone development (see discussion) (Figure 3E and F).

The close agreement between germline mutants and FØ[*foxq2*] and FØ[*nr2e3*] demonstrates that our approach is reliable (phenotypes are clear and quantifiable), flexible (mutations were created using any relevant combination of transgenic lines) and efficient in terms of cost and labor (a gene can be evaluated in less than a month by a single person, without significantly increasing burden in animal care). This motivated us to screen new and poorly characterized candidate genes with differential expression across photoreceptors, including *skor1a, sall1a, lrrfip1a* and *xbp1*.

Across multiple photoreceptor transcriptomic datasets, including ours, the expression of *skor1a* is restricted to UV and S cones (Figure 3 – figure supplement 2A) (30, 31). In humans, MEIS1 regulates expression of *SKOR1* (58). MEIS1 is key for the proper regulation of retinal progenitors across vertebrates (39, 40), making Skor1a a candidate factor that could be involved in the specification of UV and S cones (31). In disagreement with this hypothesis, we find that FØ[*skor1a*] have normal UV and S cone densities (Figure 3 – figure supplement 2B). The cone-specific gene *sall1a* is hypothesized to be involved in rod *vs*. cone differentiation in chicken (14, 53, 59). Yet, we found that FØ[*sall1a*] have normal rod densities and no disturbance of the cone mosaic (Figure 3 – figure supplement 3). Similarly, FØ[*lrrfip1a*] that have a disruption in the M-cone enriched gene *lrrfip1a*, do not produce changes in M-cone densities (Figure 3 – figure supplement 4), and FØ[*xbp1*] that have a disruption in the UV-cone enriched gene *xbp1*, have no appreciable changes in UV-cone or other photoreceptor-subtype densities (Figure 3 – figure supplement 5). These results suggest that these 4 transcription factors are not critical for photoreceptor-subtype specification. They may play other subtype-specific roles beyond specification that warrant further investigations.

### Tbx2 is a master regulator of photoreceptor fate

#### Tbx2a and Tbx2b are independently required for UV-cone specification

To further expand our FØ analysis, we explored the role of Tbx2 in photoreceptor differentiation. Tbx2 is known to be differentially expressed in cones of many species, including cichlids (60), chickens (61), squirrels (29) and primates (33). As a teleost duplicated gene, there are two paralogues of *tbx2* in the zebrafish genome: *tbx2a* and *tbx2b*. Work in zebrafish has shown that Tbx2b is involved in the determination of UV-cone fate (19). Our RNA-seq data revealed that both *tbx2a* and *tbx2b* show high expression in UV cones (Figure 4A). In addition, we detected significant enrichment of *tbx2a* and *tbx2b* expression in L and S cones respectively. This expression data suggested that Tbx2 might play numerous unexplored roles in photoreceptor specification.

**Figure. 4.**
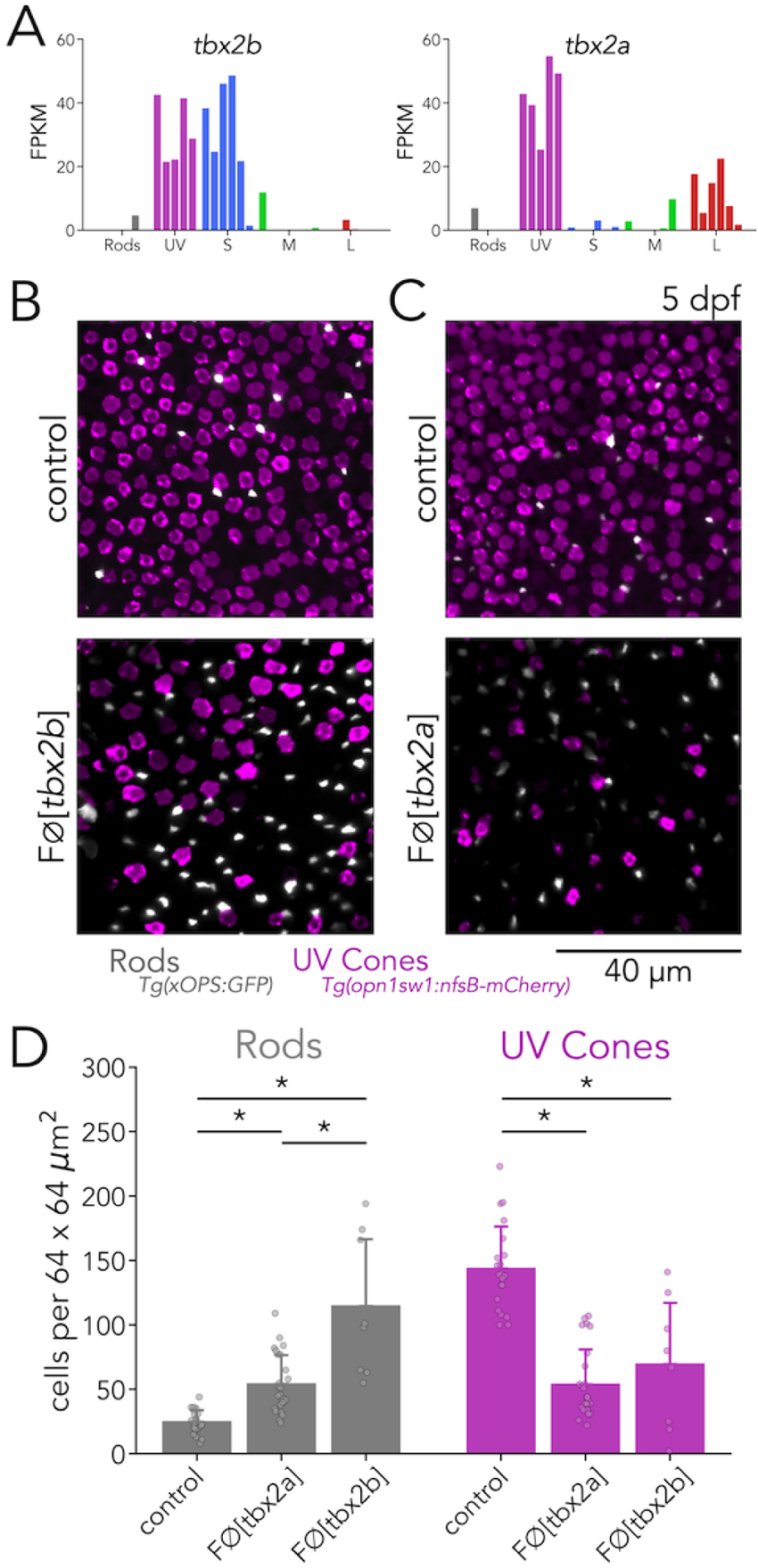
Tbx2a and Tbx2b are independently required for UV-cone specification. **(A)** *t bx2b* i s e xpressed b y b oth U V a nd S c ones (left), w hile *t bx2a* is expressed by both UV and L cones (right). **(B)** Mutations in *tbx2b* cause a loss of UV cones and an increase in rods. Representative confocal images of the central retina of control and FØ[*tbx2b*] at 5 dpf, in double transgenic larvae with labeled UV cones (magenta) and rods (grey). **(C)** Mutations in *tbx2a* also cause a loss of UV cones and an increase in rods. Representative confocal images of the central retina of control and FØ[*tbx2a*] at 5 dpf, in the same double transgenic lines. **(D)** Quantification o f r ods a nd U V c ones i n c ontrol, F Ø[*tbx2b*] a nd F Ø[*tbx2a*] larvae. Bars represent averages, error bars correspond to standard deviations, and markers correspond to individual retinas. In FØ[*tbx2b*], compared to control (n_wt_ = 21), there is a significant 2.2-fold increase in rods and a marked ∼52% decrease in UV cones (Mann-Whitney U_UV_ = 150.00, p_UV_ = 0.001, U_Rods_ = 0.00, p_Rods_ = 4.55 × 10^−5^, n_FØ[*tbx2b*]_= 8). In FØ[*tbx2a*], there is a significant ∼ 2.2-fold i ncrease i n r ods a nd a ∼63% decrease in UV cones (Mann-Whitney U_UV_ = 517.00, p_UV_ = 2.1 × 10^−8^, U_Rods_ = 44.50, p_Rods_ = 1.60 × 10^−6^, n_FØ[*tbx2a*]_= 25). The increase in rods is significantly higher in FØ[*tbx2b*] compared to FØ[*tbx2a*], while the loss of UV cones is not signoifiocantly different (Mann-Whitney U _Rods_ = 173.00, p _Rods_ = 0.002, U _UV_ = 109.00, p_UV_ = 0.72, n_FØ[*tbx2b*]_ = 8, n_FØ[*tbx2a*]_ = 25).

We first focused our analysis on the role of Tbx2 in UV cones. For our FØ analysis, we designed 3 gRNAs targeting exon 3 of *tbx2b* or *tbx2a*. In both genes, exon 3 contains critical DNA-binding residues that are completely conserved across vertebrates (62). In control larvae at 5 dpf, UV cones are numerous and densely distributed across the retina, while overall rod density is low, with most rods concentrated in the ventral retina and the lowest density in the central retina (19, 63, 64). In agreement with previous studies, FØ[*tbx2b*] had a marked decrease in UV cones (∼52%) and an increase in rod density (∼4.6 fold) (Figure 4B and D) (19). After replicating the described phenotypes of germline *tbx2b* mutants in FØ[*tbx2b*], we examined FØ[*tbx2a*]. Surprisingly, we found that FØ[*tbx2a*] displayed the same phenotype as FØ[*tbx2b*]: a marked loss of UV cones (∼63%) and an increase in rods (∼2.2 fold)—although this increase in rods was lower in FØ[*tbx2a*] than in FØ[*tbx2b*] (Figure 4C and D).

To confirm the phenotypes of *tbx2* mutants revealed through imaging of reporter lines, we quantified opsin expression using real-time quantitative PCR (qPCR). We found that, in comparison to controls, FØ[*tbx2b*] showed a clear decrease in UV-opsin expression and a significant increase in rhodopsin expression. FØ[*tbx2a*] also showed a clear decrease in UV-opsin expression, but without significant changes in rhodopsin expression (Figure 4 – figure supplement 1). Together our reporter lines and qPCR analyses suggest that, despite 87% protein-sequence similarity and co-expression of the two genes in the same cell, both Tbx2a and Tbx2b are required for the specification of zebrafish UV cones. Loss-of-function of either gene leads to a decrease in UV cones and a concomitant routing of photoreceptor progenitors towards a rod fate. In FØ[*tbx2b*] routing towards a rod fate appears to be stronger than in FØ[*tbx2a*].

#### Tbx2a inhibits M-opsin expression in L cones

After ascertaining the requirement of Tbx2a and Tbx2b in UV-cone specification, we examined whether either of these transcription factors impacted the specification of other photoreceptor subtypes. In addition to expression in UV cones, we detected significant enrichment of *tbx2a* in L cones, albeit with expression levels lower than in UV cones (Figure 4B). Furthermore, our qPCR quantification of opsins revealed a significant increase in M-opsin expression in FØ[*tbx2a*]— specifically of *opn1mws2*—along with a small decrease in L-opsin expression—specifically of *opn1lw2* (Figure 3 – Figure Supplement 1). Based on these results, we tested whether Tbx2a is involved in M-cone or L-cone specification.

To examine M and L cones, we assessed FØ[*tbx2a*] using an M-cone reporter line, where GFP expression is under direct control of the M-opsin promoter— *Tg(opn1mws2:GFP*)—in combination with an L-cone reporter line—*Tg*(*thrb:tdTomato*). In control larvae, the expression of GFP and tdTomato is non-overlapping, reflecting the distinct fate of M cones and L cones (Figure 5A, left). In FØ[*tbx2a*], we found no significant changes in the number of L cones (identified by their tdTomato expression) (Figure 5B), but there was a marked increase in the number of GFP-positive cells (presumptive M cones). Interestingly, in FØ[*tbx2a*], many GFP-positive cells co-express tdTomato (L-cone marker)—a phenotype which is not present in control larvae (Figure 5A, middle). To quantify this effect, we calculated the fraction of tdTomato-positive L cones with significant GFP expression (see methods). We found that only a small fraction of L cones are double positive in controls (mean ± s.d.: 5.2% ± 6.0), but a significantly higher fraction is double positive in FØ[*tbx2a*] (mean ± s.d.: 37.5% ± 18.9%, p < 0.01) (Figure 5C). This abnormal expression of GFP in L cones in FØ[*tbx2a*], combined with the increase in M-opsin expression found in our qPCR analysis, indicates a loss of inhibitory control over the M-opsin promoter. As a control, we repeated this M-cone and L-cone assessment in FØ[*tbx2b*]—despite no detectable expression of *tbx2b* in M or L cones (Figure 3B). While FØ[*tbx2b*] also had a significant increase in the number of GFP-positive cells (see next section) (Figure 5A, bottom), there were no changes in the number of tdTomato-positive L cones (Figure 5B) or in the fraction of L cones with significant GFP expression (mean ± s.d.: 2.6% ± 1.8%, p = 0.27) (Figure 5C).

**Figure. 5.**
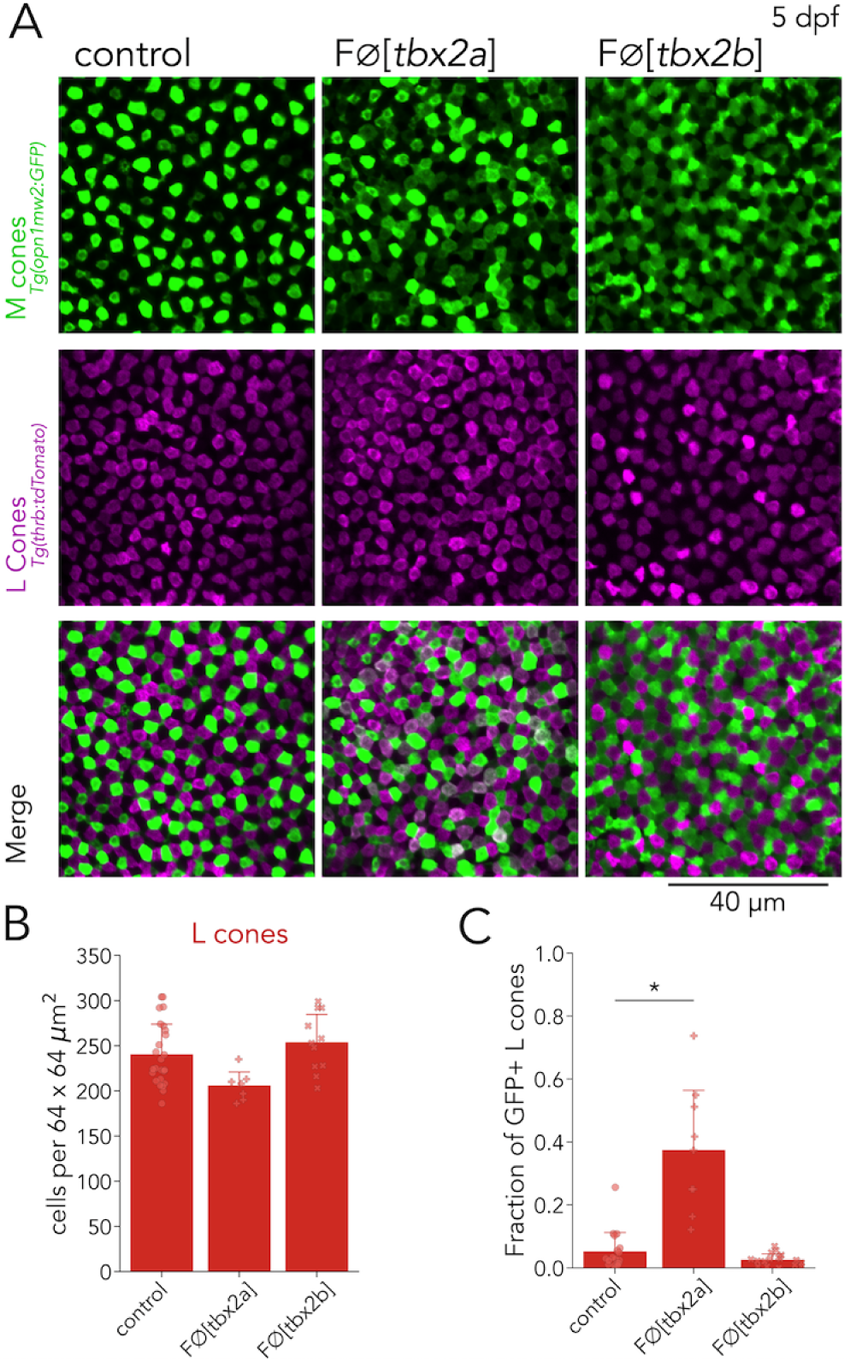
**(A)** Representative confocal images of the central retina of control, FØ[*tbx2a*] and FØ[*tbx2b*] at 5 dpf, in double transgenic larvae that label M cones— or M-opsin expressing cells—with GFP (green) and L cones with tdTomato (magenta). FØ[*tbx2a*] and FØ[*tbx2b*] display an increase in GFP-positive cells. In FØ[*tbx2a*], increase in GFP signal is restricted to tdTomato-positive cells which appear as double-positive (white) in merged images, while in FØ[*tbx2b*], increase in GFP signal is excluded from tdTomato-positive cells, producing a decrease in the space without fluorescence. **(B)** Quantification of L cones in the central retina shows no significant changes in FØ[*tbx2a*] (n_FØ[*tbx2a*]_=7) compared to control (n_wt_ = 25, Mann-Whitney U = 143, p = 0.012), or in FØ[*tbx2b*] (n_FØ[*tbx2b*]_=11, Mann-Whitney U = 104.5, p = 0.26). **(C)** Quantification of the fraction of GFP-positive L cones (double positive cells in A) reveals a significant increase only in FØ[*tbx2a*] (Mann-Whitney U = 4, p = 8 × 10^−5^), and not in FØ[*tbx2b*] (Mann-Whitney U = 186.5, p = 0.27)

These results suggest that Tbx2a, but not Tbx2b, is important to preserve L cone identity. Without Tbx2a, L cones are unable to suppress M-opsin expression (60). Overall, analysis of FØ[*tbx2a*] revealed that Tbx2a is important for UV-cone specification and for maintaining L-cone identity.

#### Tbx2b inhibits M-opsin expression in S cones

After identifying an additional role for Tbx2a, we turned our analysis to Tbx2b. Our RNA-seq revealed that in addition to UV cones, *tbx2b* is expressed in S cones (Figure 4A). Furthermore, our qPCR quantification also showed an increase in M-opsin expression in FØ[*tbx2b*]—specifically of *opn1mw1* and *opn1mw2* (Figure 4 – figure supplement 1)—and in Sopsin expression. Based on these results, we tested whether Tbx2b is involved in S- or M-cone specification.

For our analysis of S and M cones in FØ[*tbx2b*] larvae, we again used the M-opsin reporter line— *Tg(opn1mws2:GFP*)—now in combination with an S-cone reporter line—*Tg(opn1sw2:nfsB-mCherry)*. In control larvae, expression of the reporter proteins is largely nonoverlapping, except for a small fraction of S cones that consistently expresses GFP (Figure 6A, top) (28). In FØ[*tbx2b*], we did not find significant changes in the number of S cones (identified by mCherry expression) (Figure 6B), but as described above, we did observe a clear increase in the number of GFP-positive cells (presumptive M cones). Furthermore, in FØ[*tbx2b*], we found that this increase in GFP expression was restricted to S cones, which become double-positive for GFP and mCherry expression (Figure 6A, bottom). We quantified the fraction of mCherry-positive S cones with significant GFP expression, and found that, in control larvae, this fraction is low (mean ± s.d.: 9.2% ± 10.2%). In comparison, in FØ[*tbx2b*] this fraction is significantly higher (mean ± s.d.: 57.6% ± 32.2%, p < 0.01) (Figure 6C). This abnormal increase in GFP expression in S cones in FØ[*tbx2b*], combined with the increase in M-opsin expression found in our qPCR analysis, indicates a loss of inhibitory control over the Mopsin promoter. As a control, we repeated this S- and M-cone assessment in FØ[*tbx2a*]. We again observed an increase in GFP-positive cells in FØ[*tbx2a*] but without any significant changes in the number of S cones (Figure 6B) or in the fraction of mCherry-positive S cones with significant GFP expression (mean ± s.d.: 8.3% ± 4.9, p = 0.81) (Figure 6C). These results again corroborate our RNA-seq data showing that Tbx2a is not expressed by S cones and is not involved in S-cone determination.

**Figure. 6.**
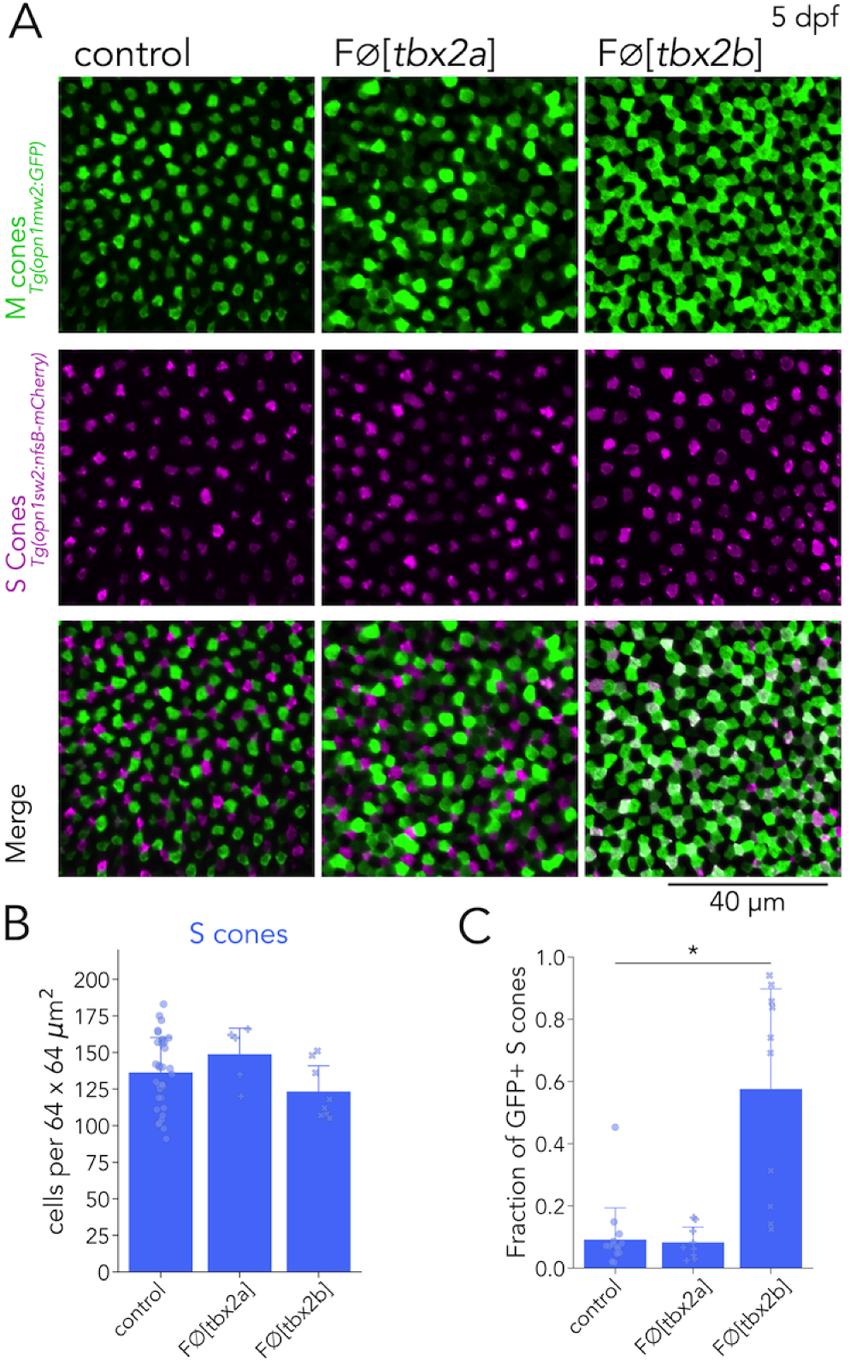
Tbx2b inhibits M-opsin expression in S cones. **(A)** Representative confocal images of the central retina of control, FØ[*tbx2a*] and FØ[*tbx2b*] at 5 dpf, in double transgenic larvae that label M cones—or M-opsin expressing cells— with GFP (green) and S cones with mCherry (magenta). FØ[*tbx2a*] and FØ[*tbx2b*] display an increase in GFP-positive cells. In FØ[*tbx2a*], increase in GFP signal is excluded from mCherry-positive cells, producing a decrease in the space without fluorescence, while in FØ[*tbx2b*], increase in GFP signal is restricted to mCherry-positive cells, which appear as double positive (white) in the merged images. **(B)** Quantification of S cones in the central retina shows no significant changes in either FØ[*tbx2a*] (n_FØ[*tbx2a*]_=5, Mann-Whitney U = 55, p = 0.21) or FØ[*tbx2b*] (n_FØ[*tbx2b*]_=8, Mann-Whitney U = 179, p = 0.17) compared to control (n_wt_ = 34). **(C)** Quantification of the fraction of GFP-positive S cones (double positive cells in A) reveals a significant increase only in FØ[*tbx2b*] (Mann-Whitney U = 6, p = 14 × 10^−5^), and not in FØ[*tbx2a*] (Mann-Whitney U = 63, p = 0.81).

In summary, further analysis of FØ[*tbx2b*] revealed that Tbx2b helps to maintain S-cone identity. Without Tbx2b, S cones are unable to suppress the expression of M-opsin. Overall, analysis of FØ[*tbx2b*] revealed that Tbx2b is important for UV-cone specification and for maintaining S-cone identity. Together with our Tbx2a results, this suggests that Tbx2 paralogs are required for the specification of UV cones and critical for establishing the distinct identities of multiple photoreceptor subtypes (Figure 7). The discovery of these new roles of Tbx2 in photoreceptor specification demonstrates the power of the methods and techniques presented in this study.

**Figure. 7.**
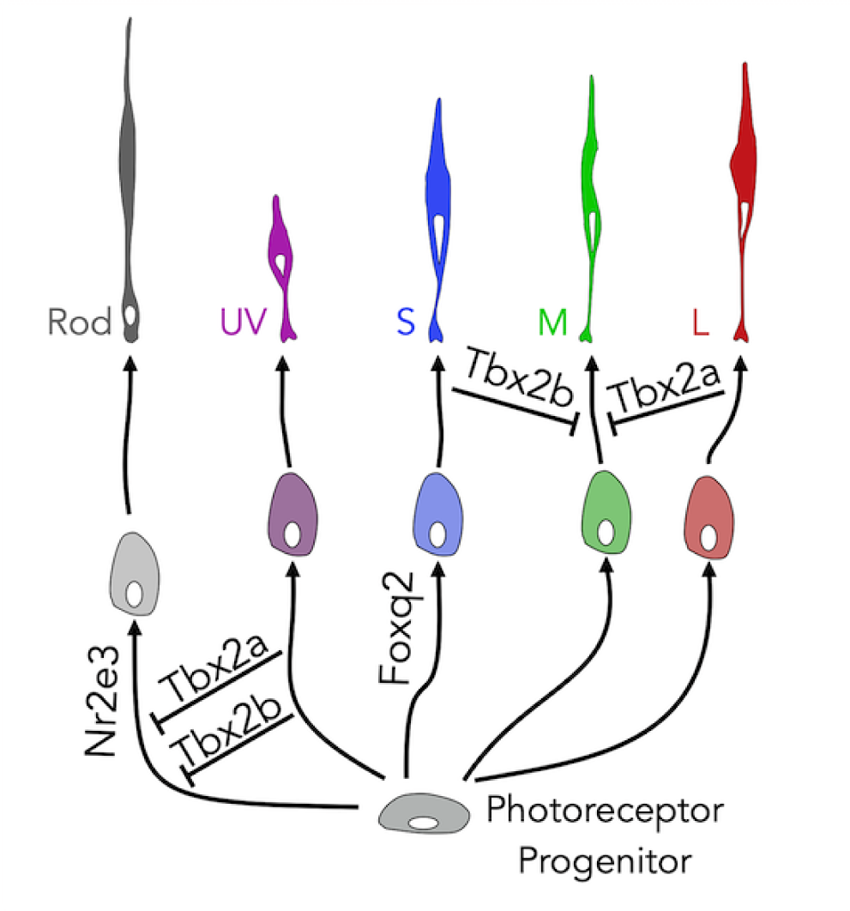
Summary diagram. The specification of zebrafish UV cones requires an early decision in photoreceptor progenitors in which Tbx2a and Tbx2b are both required to repress rod fate. The specification of zebrafish S cones depends an early decision in which Foxq2 is required to activate S-cone fate. Similarly, the specification of rods depends on Nr2e3. After their initial specification, S cones require Tbx2b to repress M-cone fate, while L cones require Tbx2a to repress M-cone fate.

## Discussion

The tools, resources and methods presented here provide a path to accelerate discovery in photoreceptor and retinal biology. We have generated transcriptomic profiles from photoreceptors with unmatched depth and purity. These transcriptomes can be used to explore previously unrecognized geneexpression patterns across photoreceptor subtypes. Importantly, we reliably identify transcription factors that may play a central role in controlling fate decisions during photoreceptor development. We also demonstrate how FØ screening can be applied as a rapid, efficient, and flexible platform to create and study loss-of-function mutations. In our study, we apply FØ-screening to investigate eight different transcription factors and test their involvement in photoreceptor specification. Together these methods provide an excellent *in vivo* setting to discover the function of other novel genes identified in our RNA-seq dataset.

### Relation to other transcriptomic datasets

Recent studies have derived transcriptomes from zebrafish retinal cells and contain information from adult photoreceptors that provide an excellent resource to benchmark the quality of transcriptomes presented here. In our study, we derived samples using manual collection for a cell-type specific, SMART-seq2-based approach (29). Three other recent studies used a variety of methods to segregate cell types in the retina. Rod transcriptomes were obtained by fluorescent-activated cell sorting (FACS) (Sun *et al*., 2018) (65). In another study, retinal-cell transcriptomes were obtained using a single-cell droplet-based (dropSeq) approach in adults and at several time points during development (Hoang *et al*., 2020) (30). Finally, transcriptomes from adult zebrafish photoreceptors were obtained by enrichment through FACS followed by dropSeq (Ogawa and Corbo, 2021) (31).

We find that there are general consistencies across these datasets, which can be exemplified by focusing on phototransduction genes: we identify rod-enrichment in 26 of 27 phototransduction genes that are known to be rod-specific, while Sun *et al*., 2018 identify 22 and Ogawa and Corbo, 2021 identify 23. We identify cone-enrichment in 31 of 35 phototransduction genes known to be cone specific, with high similarity to the subtype-specific expression patterns of Ogawa and Corbo, 2021. We found that these expression patterns are obscured in Hoang *et al*., 2020 due to contamination with rod transcripts in all the retinal cells derived from adults—many known rod-specific genes are present in all photoreceptor subtypes (Figure 1 – figure supplement 3A). Rods are the predominant cell type in the zebrafish adult retina—constituting ∼40% of all photoreceptors (25). In our experience, rods are fragile during dissociation and rod contamination presents a challenge to obtaining pure, subtype-specific datasets. Rhodopsin (*rho*) detection in non-rod samples is a simple way to assess contamination. We find that samples in Sun *et al*., 2018 and in Hoang *et al*., 2019 have significant rod contamination (> 15%), while in Ogawa and Corbo, 2021 and in the data presented here, the rod contamination is low (< 5%) (Figure 1 – figure supplement 3B). Transcriptome depth was considerably higher in our study compared to all other datasets. (Figure 1 – figure supplement 3C). The high signal-to-noise ratio in our transcriptomes allows the detection of significantly more differentially-expressed genes (DEGs). In Ogawa and Corbo, 2021, the authors detect 805 DEGs between photoreceptor subtypes (their report of ∼1100 DEGs includes those that differentiate bipolar cells from photoreceptors). In our dataset, with more stringent criteria, we identify 3058 unique DEGs (Supplementary Data 1); 598 genes are shared by both datasets, 207 are unique to Ogawa and Corbo, 2021, and 2460 are unique to this study. This higher signal-to-noise ratio is apparent for the targets of our FØ-screen—*nr2e3, foxq2, skor1a, sall1a, lrrfip1a, xbp1, tbx2a*, and *tbx2b*. In particular, the restricted expression of *tbx2a* in UV and L cones—confirmed by our FØ-screen results—is only apparent in our dataset (Figure 1 – figure supplement 3D).

Overall, we find that the methods presented in this study are especially useful to generate high-quality transcriptomes of targeted cells. High depth and low contamination increase the statistical confidence and allow the detection of genes expressed at relatively low levels (*e*.*g*., *tbx2a* expression in L cones). Our method nicely complements dropSeq approaches that sample many more cells, which is especially advantageous for discovering new cell types or tracking developmental trajectories. In our view, these techniques are complementary and integration across datasets is imperative. To facilitate such comparisons, we have created an interactive plotter that integrates analysis across the datasets as outlined here. This resource is openly available and allows easy exploration and direct comparisons across datasets (https://github.com/angueyraNIH/drRNAseq/), and includes the code and data needed to replicate our analyses. The expression plots presented here for all studies can be directly generated in this interactive plotter.

### Reliability and efficiency of FØ screening

The generation of loss-of-function mutants remains a cornerstone to test gene function. Nevertheless, creating germline mutants for all the candidate genes identified by RNAseq would require excessive effort and resources, especially given the need to make crosses into each gene of the relevant reporter lines. In our study, we use an FØ screen to accelerate the discovery of genes involved in the specification of photoreceptor subtypes. Overall, we were able to create mutations in the targeted gene in more than 80% of injected larvae by using 2 – 3 gRNAs (Table 3), and we were able to reliably phenocopy germline mutants, as exemplified by the loss of S cones in FØ[*foxq2*] and the loss of rods in FØ[*nr2e3*]. Interestingly, we find that Ø[*nr2e3*] also have a decrease in UV-cone density. We speculate that Nr2e3, which is expressed transiently by early cone progenitors (30, 66, 67), may play a role in the survival of developing UV cones, which we will pursue in the future. These findings highlight the flexibility of this screening method. In particular, the ability to evaluate the effect of gene mutations in less than a month, in any genetic background—transgenic lines or other mutant lines to investigate genetic pathways and networks—makes FØ screening a versatile technique to identify other transcription factors that may be involved in photoreceptor development and subtype specification.

### Tbx2 is a master regulator of photoreceptor fate

After the success of uncovering phenotypes in FØ[*foxq2*] and FØ[*nr2e3*], we explored the effects of *tbx2* mutations in photoreceptor identity. Our analyses revealed that Tbx2 is connected to the fate of all photoreceptor subtypes in zebrafish (Figure 7).

First, we showed that *tbx2a* and *tbx2b* are both expressed in UV cones, and the loss of either gene impairs UV-cone specification. The high conservation in the amino acid sequence of TBX2 across vertebrates and the specific expression in evolutionarily related cone subtypes (*opn1sw1*-expressing photoreceptors in zebrafish, chicken, squirrels and primates) (29, 33, 61) suggests that TBX2 may play a similar role across vertebrate species. We find that loss of UV cones in either FØ[*tbx2a*] or FØ[*tbx2a*] is associated with an increase in the number of rods during development. The switch in fate from UV cone to rod suggests that Tbx2a and Tbx2b play a role in an early fate decision in photoreceptor progenitors, allowing the acquisition of UV-cone identity by actively repressing rod fate. Interestingly the increase in rods (or rhodopsin expression) was not equal between FØ[*tbx2a*] and FØ[*tbx2b*], suggesting that the two transcription factors regulate downstream targets differently. In addition, *in vitro* experiments have shown that Tbx2 binds to DNA as a monomer (62), which makes the possibility of Tbx2a/Tbx2b dimers unlikely. Currently, it remains unclear why the specification of UV cones in zebrafish would require a “two-factor authentication” system that relies on two highly homologous but independent transcription factors.

Second, we show that *tbx2a* and *tbx2b* are expressed in L cones and S cones, respectively. Further, Tbx2a and Tbx2b help maintain L-cone and S-cone identity by repressing the expression of M opsin *in vivo*. A recent study in cichlids demonstrated that Tbx2a can bind and directly regulate the M-opsin promoter *in vitro* (60). This work also found that expression of *tbx2a* correlated strongly with the relative expression of M and L opsins, which cichlid species use to adjust their overall spectral sensitivity and match the requirements imposed by their habitats. These findings highlight that Tbx2 not only plays a role in UV specification, but is also important to maintain the identity of L and S cones.

### Outlook

While conducting the experiments described in this paper, we learned a few lessons worth highlighting. First, we find that manual picking targeted cell types allowed us to focus on collecting healthy cells and generate transcriptomes of high depth and quality. An important advantage of this method is that barriers imposed by a cell type with a low density can be largely ignored if the targeted cell types can be recognized. For this reason, we think it would be interesting to apply this technique to fully understand further subdivisions of each photoreceptor subtype including the differences between *opn1lw1*- and *opn1lw2*-expressing L cones (68) or between *opn1mw4*-expressing M cones and other M cones in zebrafish (31). Furthermore, it would be useful to explore regional specializations across the retina like the one proposed for UV cones in the acute zone (69), and for fovea *vs*. periphery differences in primates (33). This manual-picking technique is likely to also be useful beyond photoreceptors to dissect differences between subtypes of other retinal cells. Second, we find it is critical to create fast and easy access to multiple transcriptomic datasets. Eliminating technological barriers is important to ensure data can be accessed by all users. By ensuring proper access, new hypotheses pertaining to factors involved in photoreceptor development and other aspects of photoreceptor biology can be more readily explored. For example, many orthologs of human genes associated with retinal degenerations show high expression in zebrafish photoreceptors. For these reasons, we have taken a special effort to provide an interactive plotter that allows open exploration of four RNAseq datasets in a single place. We expect that this tool will be valuable to the scientific community.

Third, the results of our FØ-screen highlight some important features of how photoreceptors acquire their final identity. The process of specification seems to require several stages: defects in early stages can lead to a loss of subtypes (*e*.*g*. S cones in *foxq2* mutants) or to a change in identity (*e*.*g*. rods and UV cones in *tbx2* mutants), while defects in later stages can lead to specification deficits without loss of cells (*e*.*g*. misexpression of M opsin in *tbx2* mutants). In addition, while some transcription factors (like Foxq2 but also Thrb) mainly play a role in activating a particular fate, others (like Tbx2, but also Nr2e3 or Prdm1) play a role in inhibiting the fate of other cell types (36, 70). Because of its conserved sequence and expression, TBX2 may play a similar role in mammalian S cones—actively repressing the fate of rods. Such active repression is most likely a fundamental mechanism to maintain subtype identity throughout the life span of an organism. These mechanisms of cell identity echo beyond photoreceptors into the context of specification of any cell type. In fact, TBX2 plays a similar repressive role in the inner ear of mice (71).

Our current study focused on the differential expression of transcription factors because of their central role in controlling fate decisions. Moving beyond cell fate, in the future approaches like those outlined here can be used to identify and study the function of genes involved in phototransduction, metabolism, ciliary transport, synaptic machinery, *etc*. It is likely that the other targets of our FØ screen—*skor1a, sall1a, lrrfip1a* and *xbp1*—that have no clear involvement in the specification of photoreceptor subtypes, may play a role in regulating these other aspects of photoreceptor biology. The dataset and methods described here are an excellent resource to propose hypotheses, to generate an initial list of candidate genes and to perform efficient screening for phenotypes related to these other functions.

## Materials and Methods

### Animals

We grew zebrafish larvae at 28°C in E3 embryo media (5 mM NaCl, 0.17 mM KCl, 0.33 mM CaCl2, and 0.33 mM MgSO4) under a 14 h:10 h light-dark cycle. At 1 dpf, we added 0.003% 1-phenyl-2-thiourea (PTU) to the embryo medium to block melanogenesis. All work performed at the National Institutes of Health was approved by the NIH Animal Use Committee under animal study protocol #1362-13. For RNAseq samples with adult zebrafish, animals of both sexes were used. For the FØ-screen, larvae were examined at 5 dpf. At these ages, sex cannot be predicted or determined, and therefore sex of the animals was not considered. Transgenic lines used in this study are listed in Table 1.

### RNA-seq sample collection

We euthanized adult zebrafish by immersion in ice-cold water (below 4°C) followed by decapitation. We pierced the cornea with a 30-gauge needle and removed the cornea and lens before performing enucleation. Once the eye was isolated, we gently separated the retina from sclera and RPE using fine forceps or electrically-sharpened tungsten electrodes (74) and immediately started incubation in papain solution (5 U/mL papain Calbiochem#5125, 5.5 mM L-Cysteine, 1 mM EDTA in divalent-free Hank’s balanced salt solution) for 35 minutes at 28°C. After a brief wash in DMEM supplemented with 5% bovine serum albumin, we performed mechanical trituration of the retina with the tip of a 1 mL pipette and used a cell-strainer polystyrene tube to obtain a single-cell suspension. After spin-down (2000x G for 2 min), we resuspended cells in 500 µL of enzyme-free fresh DMEM and diluted the cell suspension into three serial 10-fold dilutions before plating in glass-bottom petri dishes. The dilutions ensured that we could find a preparation where the density of cells and debris was low and most photoreceptors were truly isolated. We inspected the cell suspension using an epifluorescence microscope (Invitrogen EVOS cell-imaging system) and collected 20 photoreceptors per retina based on their fluorescence and morphology (prioritizing cells that looked healthy, had intact outer segments, visible mitochondrial bundles and undamaged cell membranes) using an oil-based microinjector system (Eppendorf CellTram 4R) and glass pipettes with a 15 µm opening (Eppendorf TransferTip-ES). After collection, we resuspended photoreceptors in 1 µL of fresh PBS, reinspected cells for fluorescence, collected them in a PCR tube containing 8 µL of lysis buffer of the RNA kit and kept the tube on ice until cDNA libraries were prepared. We used the SMART-seq v4 ultra-low input RNA kit for sequencing (Takara #634897) using the manufacturer’s instructions for single-cell samples, followed by the Low Input Library Prep Kit v2 (Takara #634899). We pooled up to 12 samples (with different barcodes) in one lane of a flow cell for sequencing (Illumina HiSeq 2500) and used a 150 bp paired-end read configuration. The first sequencing batch contained 4 UV-cone and 4 S-cone samples in a single flow cell, and the second sequencing batch contained the rest of the samples divided across 2 flow cells (6 rod, 1 UV-cone, 2 S-cones, 7 M-cones and 6 L-cone samples).

### RNA-seq data analysis

After an initial quality control and trimming of primer and adapters sequences using Trimmomatic (75), we used the NIH high-performance computing resources (Biowulf) to align reads to the zebrafish genome (*Danio rerio GRCz11*) using HiSat2 (76) and to assemble and quantify transcripts using Stringtie (77). We performed differential expression analysis using Deseq2 and pcaExplorer for initial visualizations (78, 79). Genes were considered as differentially expressed if fold-enrichment> 1.5, p-value < 0.01 and the estimated false positive rate or p-adjusted < 0.1. In addition, genes were required to have positive reads in > 50% of the enriched samples. To be able to detect differences that relied on expression on just 1 or 2 cone subtypes, we removed the requirement on fold-enrichment in rod *vs*. cone comparisons. To further explore the data, we transformed read numbers into fragments per kilobase per million reads (FPKM) (Supplementary Data 01) and developed custom routines in Python for plotting. We subselected transcription factors by selecting genes identified with “DNA-binding transcription factor activity” in ZFIN (80) and repeated principal component and differential expression analyses (Supplementary Data 02 and 03). Transcription factors were considered as significantly expressed if at least 20% (*i*.*e*., 7 out of 35) of the samples had positive reads. To ensure broad access to our transcriptomic data, we provide access to the raw data (GEO accession number GSE188560), and after analysis in several formats including as a plain csv file, as a Seurat object for easy integration with dropSeq datasets (81), and finally, as an interactive database for easy browsing and visualization (). In order to make direct comparisons between our data and other RNAseq studies, we have integrated visualizations that use their publicly available data. For rod transcriptomes obtained using FACS we used the provided analyzed data, which includes gene log counts per million (cpm) for 4 rod samples (GFP-positive cells) and 4 non-rod samples (GFP-negative retinal cells) (GSE100062) (65). For transcriptomes from adult photoreceptors obtained using FACS followed by dropSeq (31), we used the Seurat object provided by the authors (GSE175929) and we used custom scripts in R (82), using Rstudio (83) and Seurat (81) to export average expression values and percent of cells with positive counts of each gene for each cluster. For transcriptomes of retinal cells obtained using dropSeq (30), we used the Seurat object for zebrafish development provided by the authors (http://bioinfo.wilmer.jhu.edu/jiewang/scRNAseq/), and we updated the object to Seurat v03 (84), extracted cells that corresponded to adult rods and cones, performed clustering and used the expression of opsins and other markers to identify cone subtypes (including *arr3a* for L and M cones, *arr3b* and *tbx2b* for UV and S cones, *thrb* and *si:busm1-57f23*.*1* for L cones and *foxq2* for S cones). All results and scripts necessary to recreate these analyses are also provided openly (https://github.com/angueyraNIH/drRNAseq).

### Quantitative PCR (qPCR)

We euthanized groups of 20 to 30 zebrafish larvae by immersion in ice-cold water (below 4°C) and immediately performed RNA extraction using the RNeasy Mini Kit (Qiagen) and reverse transcription using the High-Capacity cDNA Reverse Transcription Kit (Thermo Fisher), which relies on random primers. Samples were kept frozen (−20°C) until use. For qPCR assays we used the PowerUp SYBR Green Master Mix (Thermo Fisher) and a 96-well system (CF96, Biorad) following manufacturer’s protocols. We estimated expression levels using the relative standard curve method, using 5 serial standard dilutions of cDNA obtained from wild-type larvae. To calculate fold differences in gene expression, we normalized transcripts levels to the levels of actin-b2 (*actb2*), and all measurements were repeated in triplicate. We performed statistical testing using Mann-Whitney rank sum tests, with a p-value < 0.05 required for significance. All primers used for qPCR are provided in Table 2.

**Table 2.**
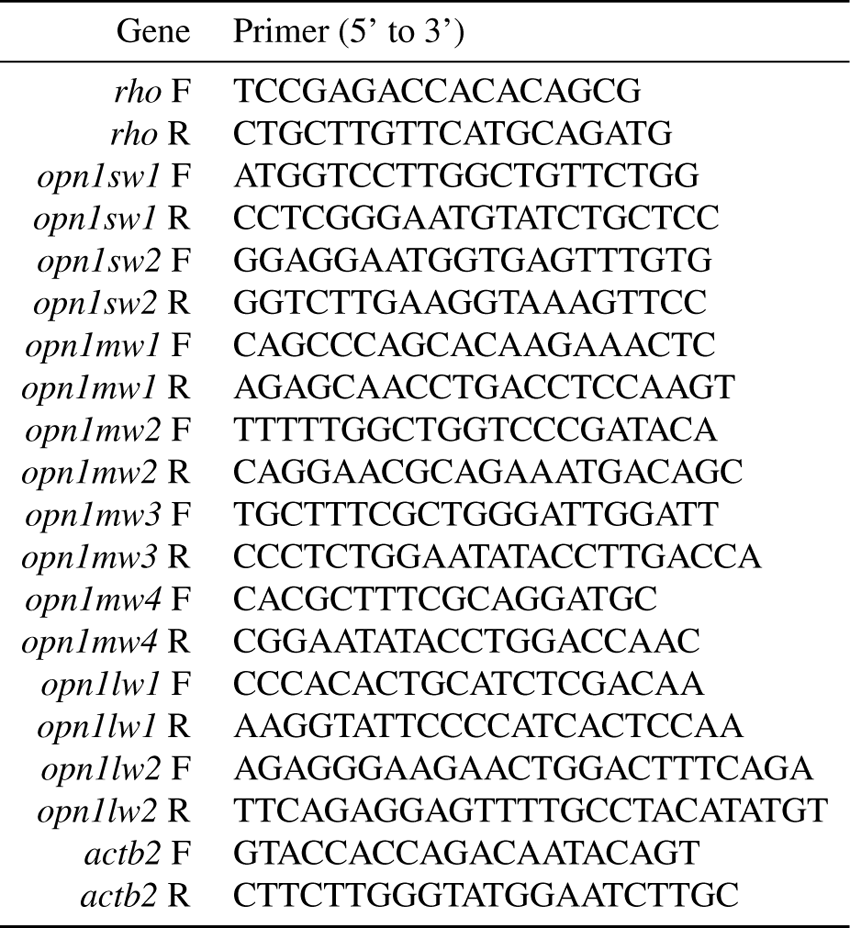
qPCR primers

### Fluorescent in situ hybridizations (RNAscope)

We performed the RNAscope assay following manufacturer’s instructions (ACDBio), with the following custom made probes: negative control (T1-T12), *actb2*-T2, *gnat2*T3, *foxq2*-T2, *tbx2a*-T3, *skor1a*-T6, *lrrfip1a*-T7, *cux2b*-T10, *smad5*-T11, *ahr1b*-T2, *etv5a*-T3. Briefly, after euthanasia, we collected eyes from adult zebrafish, embedded them in plastic molds filled with cryo-embedding medium (OCT) and froze immediately at −80°C. We obtained 15 µm cryo-sections and stored them at −80°C until use. We fixed retinal sections by immersion in 4% paraformaldehyde for 60 minutes, performed washes with RNAase-free PBST (PBS + 0.01% Tween) and dehydration in methanol in a step-wise manner (5 minutes incubation each in 25%, 50%, 75% and 100% methanol in PBST), before air drying for 5 minutes. We then applied Protease III for 5 minutes at room temperature, performed 3 washes with PBST, and hybridization with probes for 2 hours at 40°C in a humidified tray. After hybridization, we interleaved three 5-minute washes (with the provided Wash Buffer) and incubation with Amp1, Amp2, Amp3 solutions for 30 minutes, and with the Fluoro solution for 15 minutes at 40°C in a humidified tray. After the final washes, the sections were immediately covered with mounting medium and a coverslip before imaging. To combine this assay with reporter lines, we omitted the protease treatment, but this led to a decrease in staining. Decreasing Protease treatment from the recommended 30 minutes to 5 minutes improved the morphology of the tissue but did not preserve GFP fluorescence.

### FØ-CRISPR screening

We designed guide RNAs (gRNAs) using the online resource CHOPCHOP (85). We picked guides that targeted exons that encode the DNA-binding domains of transcription factors, had no self-complementarity, and that had 3 or more mismatches with other regions of the zebrafish genome; if this was not possible, we targeted the first coding exon. We used purified Cas9 protein (Alt-R® S.p. Cas9 nuclease, v.3) and chemically synthesized AltR-modified crRNA and tracrRNA (Integrated DNA technologies) for injections (22) (Table 3). We prepared 1 µL aliquots of a 25 µM stock solution of Cas9 protein diluted in 20 mM HEPES-NaOH (pH 7.5), 350 mM KCl and 20% glycerol, and stored them at −80°C until use. We diluted each target-specific crRNA and the common tracrRNA using the provided duplex buffer as a 100 µM stock solution and stored them at −20°C. We prepared a 50 µM crRNA:tracrRNA duplex solution by mixing equal volumes of the stock solutions followed by annealing in a PCR machine (95°C, 5 min; cooling 0.1°C /s to 25°C; 25°C for 5 min; rapid cooling to 4°C), then we used the duplex buffer to obtain a 25 µM stock solution, before mixing equal volumes of the guides targeted to a single gene (3 guides for *skor1a, sall1a, xbp1, tbx2a* and *tbx2b*, 2 guides for *foxq2* and *nr2e3*), making aliquots (2 µL for *foxq2* and *nr2e3*, 3 µL for the other genes) and storing at −20°C until use. Prior to microinjection, we prepared 5 µM RNP complex solutions by mixing 1 µL of 25 µM Cas9, 1 µL of 0.25% phenol red and 3 µL of the *tbx2a* or *tbx2b* duplex solution, or 1 µL of pure water and 2 µL of the duplex solution for the other genes. We incubated the RNP solution at 37°C for 5 minutes and kept at room temperature for use in the following 2 – 3 hours. We injected ∼1 nL of the 5 µM RNP complex solution into the cytoplasm of one-cell stage zebrafish embryos.

**Table 3.**
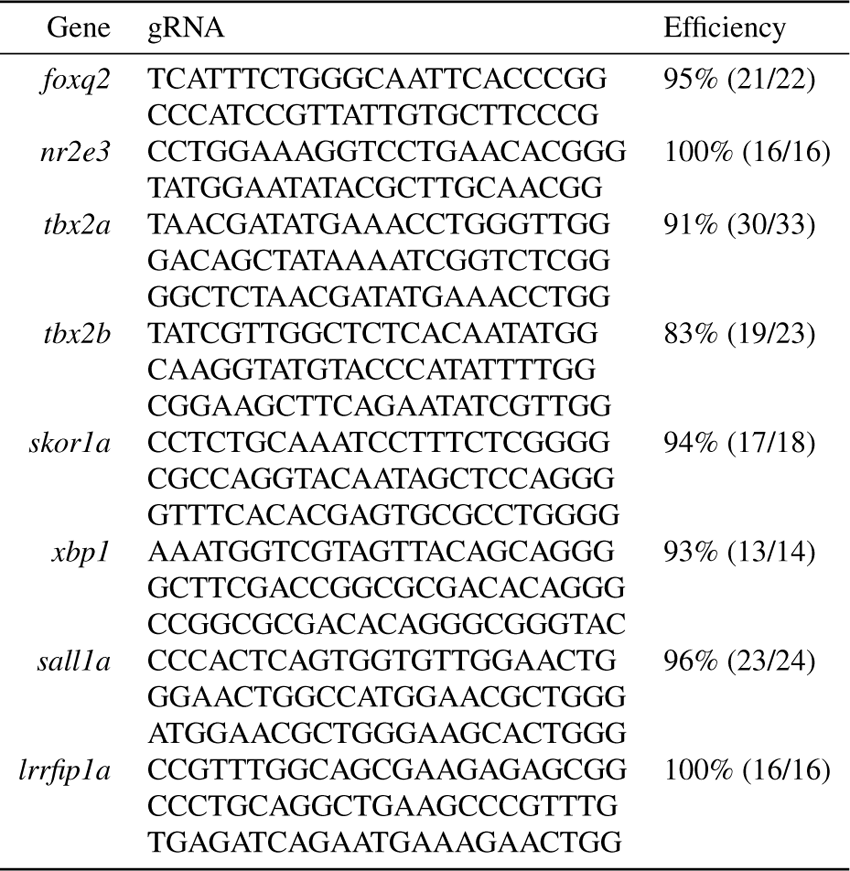
crRNA Sequences

### Genotyping

We extracted DNA from the bodies of larvae (5 dpf) after enucleation by placing them in 25 µL of 25 mM NaOH with 0.2 mM EDTA, heating to 95°C for 30 minutes, and cooling to 4°C. Then we neutralized the solution by adding 25 µL of 40 mM Tris-HCl and vortexed the samples. For genotyping, we used a fluorescent PCR method (86). We added the M13F adapter sequence (5’-TGTAAAACGACGGCCAGT-3’) to forward primers and the PIG-tail adapter sequence (5’-GTGTCTT-3’) to reverse primers and used incorporation of fluorescent M13F-6FAM for detection. Our PCR mixture (1x), for a 20 µL reaction, contained forward primer (0.158 µM), reverse primer (0.316 µM), M13-FAM (0.316 µM, IDT), Phusion HF PCR Master Mix (1x, BioLabs), water (6.42 µL), and 2 µL of DNA. We used the following PCR protocol: 1) 98°C denaturation for 30 seconds, 2) 34 cycles of 98°C for 10 seconds, 64-67°C for 20 seconds, 72°C for 20 seconds 3) final extension at 72°C for 10 minutes, 4) hold at 4°C. All primers and expected sizes are provided in Table 4, and the estimated efficiency of producing mutations with each guide combination in preliminary experiments is included in Table 3. Because of the high homology between *tbx2a* and *tbx2b*, we also tested cross-reactivity of the guides between these two genes, and found no sign of mutations in the non-targeted gene (0/48 larvae tested).

**Table 4.**
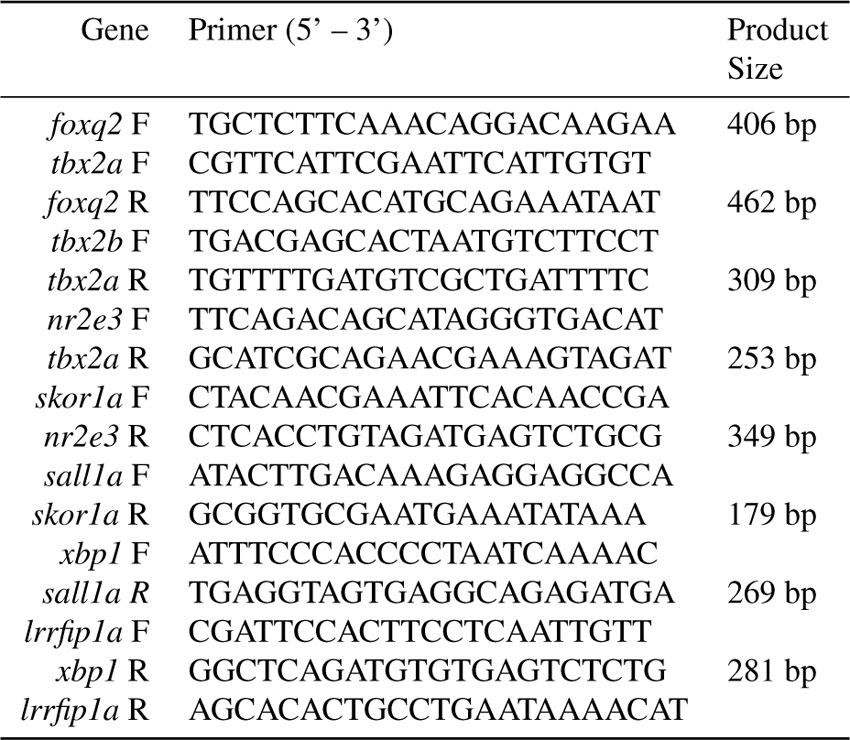
Primer Sequences

### Immunohistochemistry

We fixed zebrafish larvae at 5 dpf in 4% paraformaldehyde in phosphate buffered saline (PBS) for 1 hour at room temperature, followed by washes with 1% Triton X-100 PBS (3 × 10 min). We incubated larvae in primary antibodies diluted in 2% normal donkey serum (Jackson ImmunoResearch) and 1% Triton X-100 PBS for five days at 4°C with continuous and gentle shaking. To label S cones, we used a rabbit polyclonal anti-blue opsin (Kerafast EJH012) in a 1:200 dilution. After incubation with primary antibodies, we performed washes with 1% Triton X-100 PBS (3 × 15 min). We incubated larvae in donkey polyclonal secondary antibodies labeled with Cy5 (Jackson ImmunoResearch) in 1% Triton X-100 PBS overnight at 4°C with continuous and gentle shaking and performed washes in 1% Triton X-100 PBS (3 × 15 min) before mounting.

### Imaging

#### Sample Preparation and Image Acquisition

For *in situ* hybridizations of retinal sections, we used the same mounting medium and imaging system, but used an 60x, 1.40 NA oil-immersion objective and acquired z-stack images from a 70 µm x 70 µm square area centered on the photoreceptor layer, every 0.25 µm at a 1024 × 1024 pixel resolution. Images correspond to maximal-intensity projections of 2 – 4 µm in depth, after contrast adjustments to reject the autofluorescence of outer segments and highlight the bright fluorescent puncta.

For larval imaging, we enucleated eyes from fixed larvae using electrically sharpened tungsten wires (74). We placed isolated eyes on a coverslip and oriented eyes to place photoreceptors closest to the coverslip before using a small drop of 1.5% low-melting point agarose to fix them in place. Upon solidification, we added a polyvinyl-based mounting medium (10% polyvinyl alcohol type II, 5% glycerol 25 mM Tris buffer pH 8.7 and 0.5 µg/mL DAPI) and placed the coverslip on a glass slide, separated by a spacer (Grace Biolabs and/or duct tape) to avoid compression. We used the bodies of the larvae for genotyping and imaged the corresponding larval retinas using a Nikon A1R resonant-confocal microscope with a 25x, 1.10 NA water-immersion objective. We acquired z-stack images from a 64 µm x 64 µm square area of the central retina (dorsal to optic nerve) for photoreceptor quantification every 0.4 – 0.5 µm at a 1024 × 1024 pixel resolution.

#### Image Analysis

##### Photoreceptor quantification

We imported confocal z-stacks of the central region of the retina (64 µm x 64 µm) into Napari (87). We created maximum intensity projections (MIPs) using a small subset of the z-stack (2 – 10 planes) that ensured that we captured all photoreceptor cells in the region into a single image. We then used the Napari plugin of Cellpose, a machine-learning based segmentation algorithm, to segment photoreceptors in each image, using the *cyto2* model (88). Finally, we manually corrected the segmentation to ensure all photoreceptors were properly counted. We performed statistical comparisons for counts of each photoreceptor subtype between wild-type *(wt*) and mutant (*FØ*) larvae using Mann-Whitney rank sum tests, with a p-value < 0.01 required for statistical significance.

Identification of double-positive cells in *tbx2* mutants: The increase in the *Tg(opn1mws2:GFP)*+ cells in FØ[tbx2a] and FØ[tbx2b] larvae made segmentation of the green channel difficult and unreliable, as these additional cells did not conform to the normal spatial separation between M cones. For this reason, we exploited the more accurate segmentation of L cones and S cones using the red channel, when imaging *Tg(thrb:tdTomato)* or *Tg(opn1sw2:nfsB-mCherry)*, respectively, and used it to create masks for the green channel. We normalized the GFP signal across the whole image to span a 0 – 1 range (to be able to make comparison between images) and used a 10-pixel erosion (to avoid effects due to optical blurring during imaging of the GFP signal) before calculating the average normalized GFP signal contained within each S-cone or L-cone. By plotting the distribution of GFP signal in L cones, we were able to establish a threshold of 0.195 that was exceeded by only 5.2% of L cones in *wt* larvae and used it to classify L cones as GFP+ in both *wt* and FØ larvae. In the original work that established the *Tg(opn1mws2:GFP)* line, it was noted that a subset of S cones in *wt* larvae are GFP+ (28). We were able to identify these cells using a GFP signal threshold of 0.255 (9.2% of *wt* S cones), and again used this same threshold to quantify the fraction of GFP+ S cones in both *wt* and FØ larvae.

#### Statistical analyses

We performed statistical analyses and data plots using Python in Jupyter notebooks (89). Values of data and error bars in figures correspond to averages and standard deviations, and for statistical comparisons we used Mann-Whitney U tests. Samples sizes and significance levels are stated in the figure captions. No randomization, blinding, or masking was used for our animal studies, and all replicates are biological. For RNAseq, we performed an initial sequencing run after collecting dissociated photoreceptors in squirrel (29) and zebrafish and established that a minimum of four samples per subtype were required to establish reliable statistical significance in differential gene-expression analysis. For FØ screening, our initial experiments were aimed at replicating the loss of UV cones and the increase in rods reported for *tbx2b* mutants (19), and we established that a minimum of 6 injected larvae per group were needed to provide enough statistical power in photoreceptor quantifications in FØ larvae. We adopted p<0.01 as our value for significance for photoreceptor subtype quantifications. Injected larvae that had normal (wild-type) genotypes—a sign that CRISPR mutagenesis was not successful—were excluded from analysis, so that quantifications rely solely on larvae with confirmed mutations in the targeted gene.

## Supplemental Materials

**Supplementary data 01: Differential gene expression in zebrafish photoreceptors**. Collection of CSV files containing output of differential gene expression analysis using De-SEQ2, along relevant directions (rods *vs*. cones; (UV+S) *vs*. (M+L); M *vs*. L; UV *vs*. S) and including counts (in FPKM) for all detected genes.

**Supplementary data 02: Differential transcription factor expression in zebrafish photoreceptors: rods vs. cones**. CSV file containing transcription factors with significant differential expression between rod and cone samples.

**Supplementary data 03: Differential transcription factor expression in zebrafish photoreceptors: cone subtypes**. Collection of CSV files containing transcription factors with significant differential expression between cone subtypes.

## Data Availability

Sequencing data have been deposited in GEO under accession GSE188560. Data can be visualized at https://github.com/angueyraNIH/drRNAseq.

## Acknowledgements

This work was supported by the National Eye Institute Intramural Research Program (W.L.), the National Institute on Deafness and Other Communication Disorders Intramural Research Program (*1ZIADC000085-01*, K.S.K), and National Eye Institute Pathway to Independence Award (*K99EY030144-01*, J.A.). This work utilized the computational resources of the NIH HPC Biowulf cluster. (http://hpc.nih.gov). We would like to thank Jamie Sexton, Alisha Beirl and Katherine Pinter for all the animal care and technical support, John Ball, Elizabeth Cebul and other members of the Kindt Lab and the Li Lab for useful discussions, and Matthew Brooks, Linn Gieser and Anand Swaroop for sequencing services. We are very grateful to Rachel Wong, Takeshi Yoshimatsu, Ralph Nelson, James Fadool, Steven Leach, Brian Perkins and Xiangyun Wei for providing the transgenic zebrafish lines used in this study.

---

**Figure 1 - figure supplement 1.**
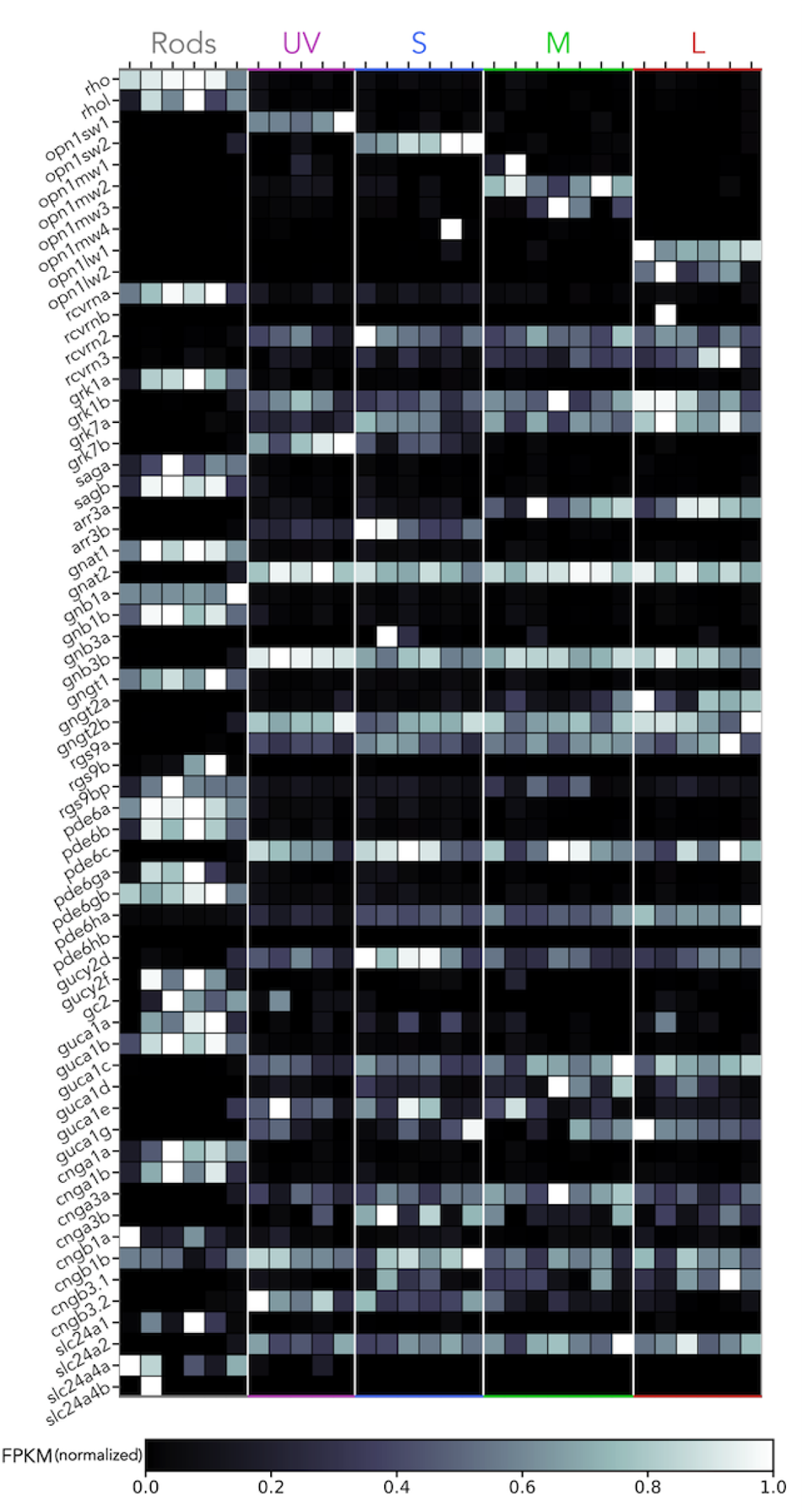
Expression of phototransduction genes. Heatmap showing expression of phototransduction genes for each RNAseq sample. Grey values indicate expression level normalized in each row by the maximal value. Genes have been arranged by functional family.

**Figure 1 - figure supplement 2.**
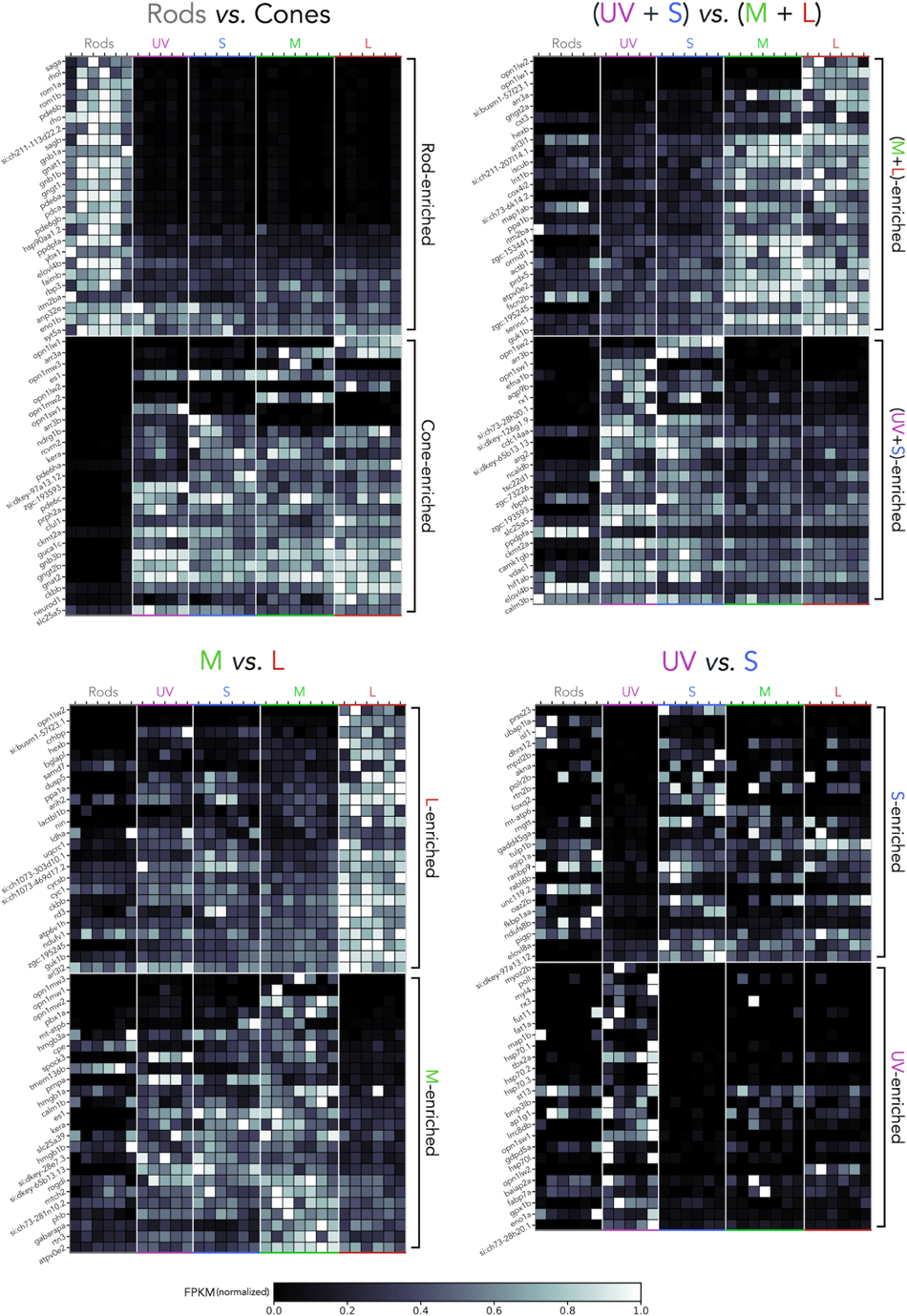
Differentially-expressed genes between photoreceptor subtypes. Heatmaps showing the top 50 differentially-expressed genes, identified using pairwise comparisons that follow the direction of principal components shown in Figure 1G, and indicated in the titles of each panel. Grey values indicate expression level normalized in each row by the maximal value. Genes have been arranged by degree of enrichment.

**Figure 1 - figure supplement 3.**
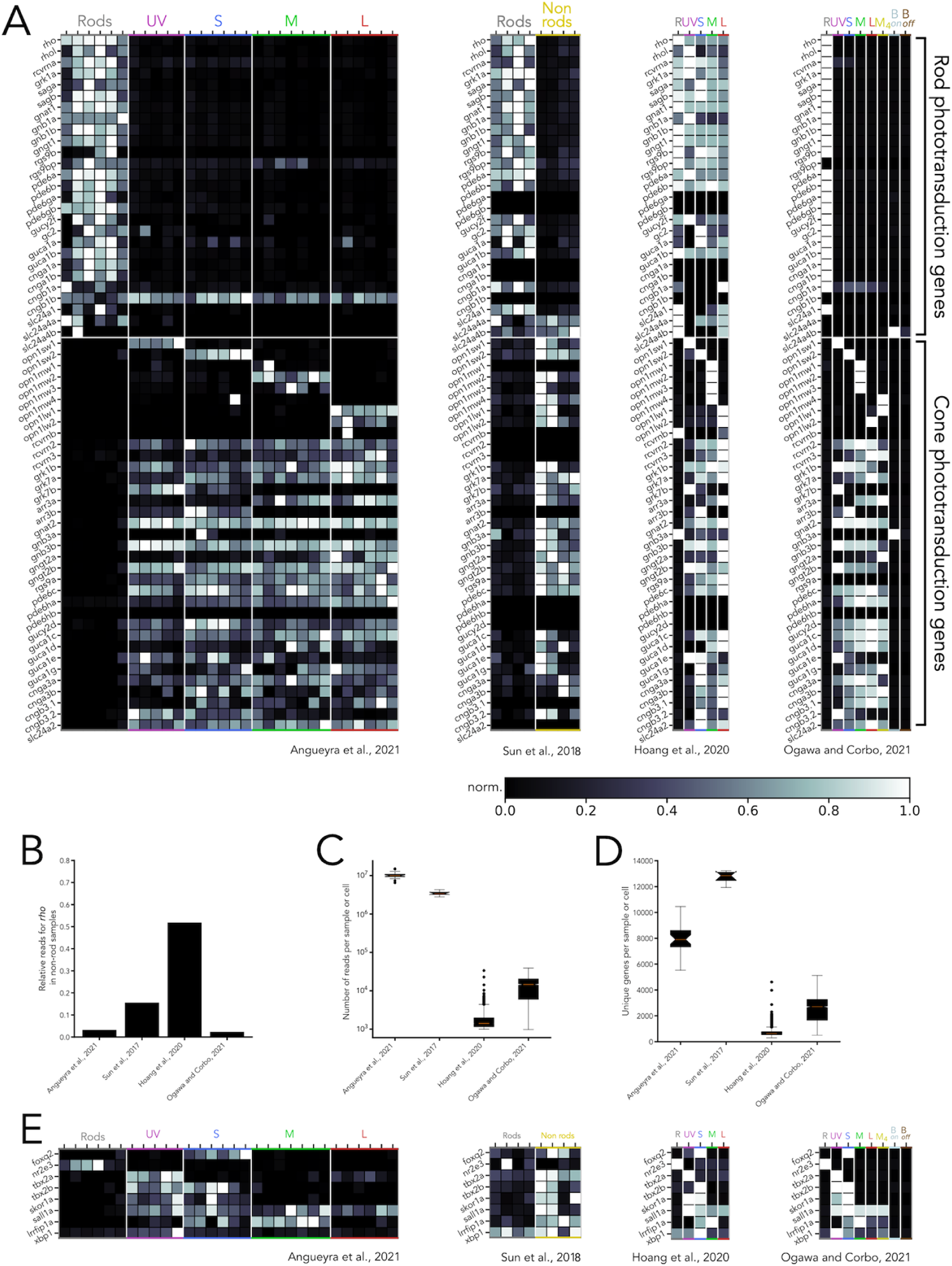
Comparison of RNAseq datasets across studies. **(A)** Heatmaps showing expression of phototransduction genes across four RNAseq studies containing transcriptomes of adult zebrafish photoreceptors. Grey values indicate expression level normalized in each row by the maximal value, and genes have been first arranged by their known expression in rods or cones and subsequently by functional family. **(B)** Degree of contamination in each study, measured as expression of rhodopsin (*rho*) in non-rod samples or cells, normalized to expression in rod samples or cells. **(C)** Transcriptome depth in each study, measured as the number of reads per sample (Angueyra *et al*., 2021, Sun *et al*., 2018), or number of reads per cell (Hoang *et al*., 2020, Ogawa *et al*., 2021), shown as a box and whiskers plot in log scale (red line corresponds to mean, boxes correspond to inter-quartile ranges, whiskers correspond to extremes and symbols correspond to outliers). **(D)** Number of unique genes per sample or per cell identified in each study. Of note, for Sun *et al*., 2018, 4 samples included all retinal cells (non rods), while for Hoang *et al*., 2020 and Ogawa *et al*., 2021, only cells clustered as photoreceptors were included. **(E)** Heatmaps showing expression of the eight transcription factors explored in this study, normalized by row. In heatmaps, R = rods, B_on_ = on bipolar cells, B_off_ = off bipolar cells.

**Figure 2 - figure supplement 1.**
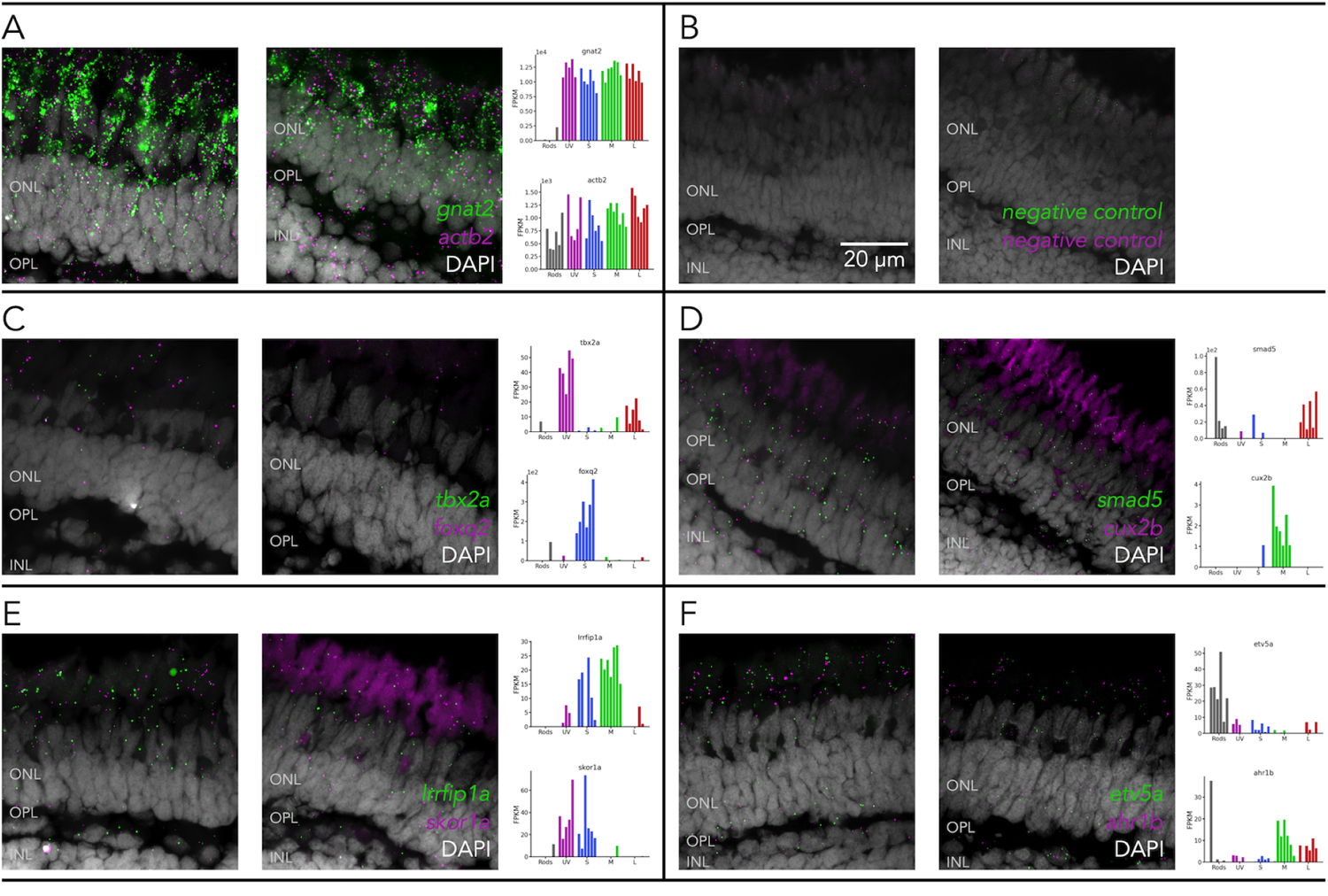
Transcripts from candidate transcription factors are detected in photoreceptorsn. Representative confocal images of fluorescent *in-situ hybridizations* (RNAscope) performed in fresh-frozen cryosections of adult zebrafish retina, counterstained with a nuclear label (DAPI, gray), to detect transcripts of **(A)** *gnat2* (green) and *actb2* (magenta), as positive controls, **(B)** of negative controls and **(C-F)** of transcription factors, identified by RNAseq as differentially expressed (plot insets), including **(C)** *tbx2a* (green) and *foxq2* (magenta), **(D)** *smad5* (green) and *cux2b* (magenta), **(E)** *lrrfip1a* (green) and *skor1a* (magenta), **(F)** *etv5a* (green) and *ahr1b* (magenta). For all probes, excluding negative controls, bright fluorescent punctae corresponding to RNA molecules for the target gene, could be detected in the photoreceptor layer. Dimmer and diffuse autofluorescence in the red channel, derived from photoreceptor outer segments, can also be seen in several of the example images. ONL = outer nuclear layer, OPL = outer plexiform layer, INL = inner nuclear layer.

**Figure 3 - figure supplement 1.**
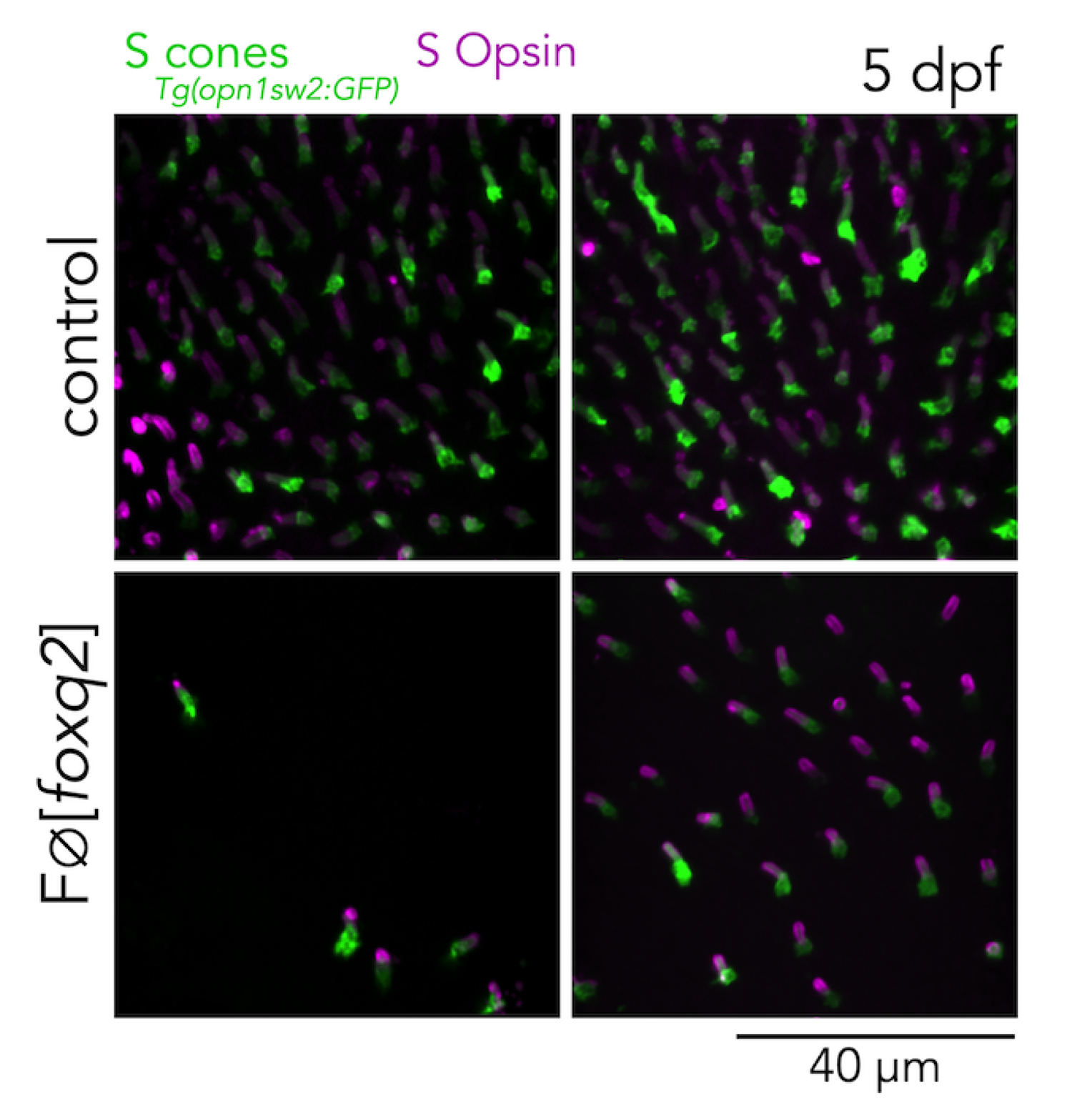
Mutations in *foxq2* cause a decrease in S opsin positive photoreceptors. Confocal images of the central retina of 2 control and 2 FØ[*foxq2*] larvae at 5 dpf, labeled with an S opsin antibody (magenta), in an S-cone reporter line (green). In the control images, all GFP-positive cells are also positive for S opsin antibody labeling (the curvature of the larval eye causes some apparent uneven labeling). In FØ[*foxq2*] larvae there is a significant loss of both GFP and S opsin antibody labeling, and remaining GFP-positive cells are also positive for S opsin.

**Figure 3 - figure supplement 2.**
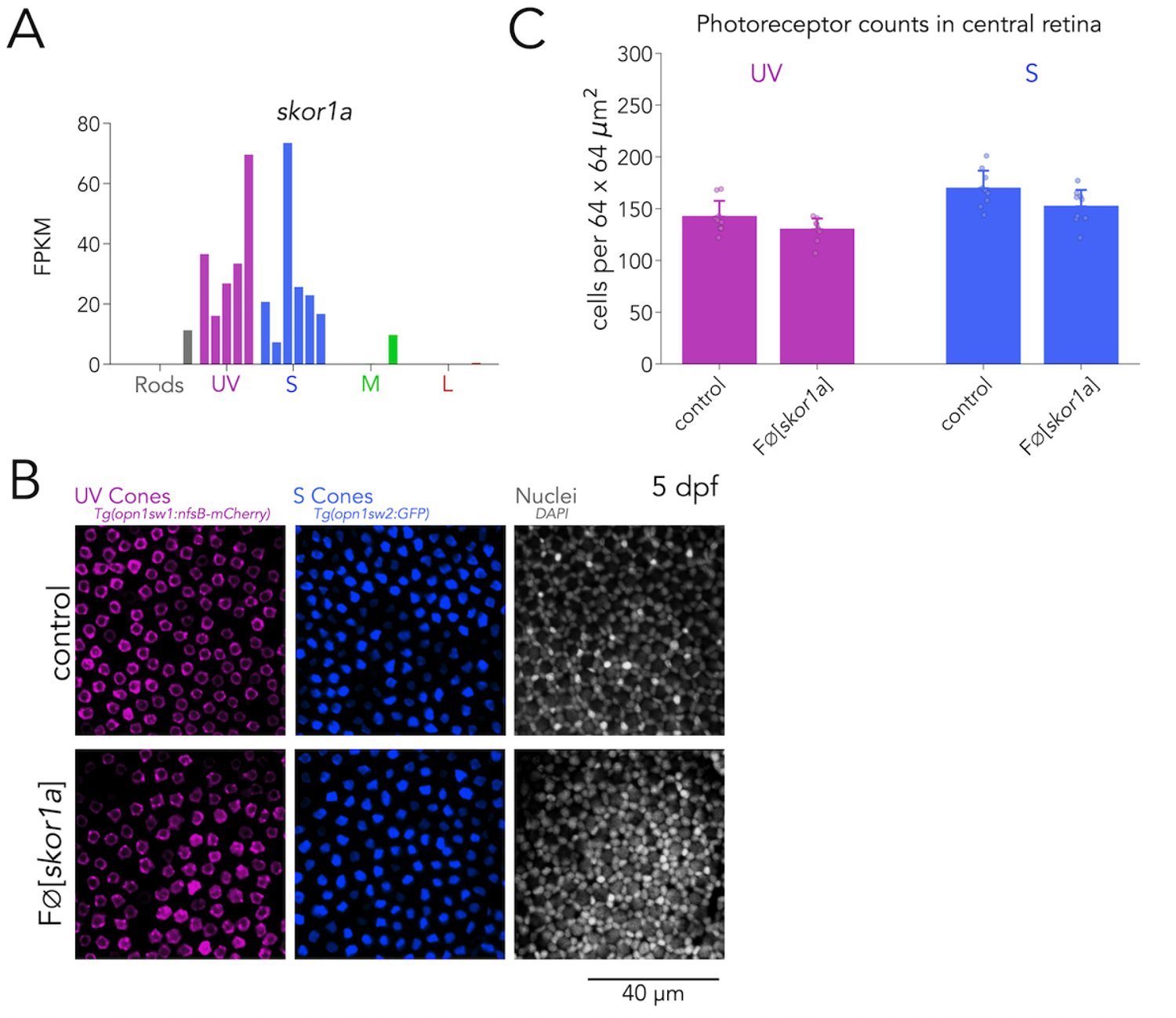
Skor1a is not required for UV-cone or S-cone specification. **(A)** Expression of *skor1a* across photoreceptors shows enrichment in UV and S cones. **(B)** Mutations in *skor1a* do not change UV or S cones and cone mosaic is not disrupted. Representative confocal images of the central retina of control (top row) and FØ[*skor1a*] (bottom row) larvae at 5 dpf. Columns correspond to transgenic line that label UV cones (magenta) or S cones (blue) or to nuclei labeled with DAPI (grey) to reveal the cone mosaic. **(C)** Quantification of photoreceptor densities in control and FØ[*skor1a*] larvae. Bars represent averages, error bars correspond to standard deviations, and markers correspond to individual retinas. There is no significant change in the density of UV cones in FØ[*skor1a*] compared to controls (Mann-Whitney U = 59.0, p = 0.11, n_wt_ = 9, n_FØ[*skor1a*]_ = 9) or of S cones in FØ[*skor1a*] compared to controls (Mann-Whitney U = 62.50, p = 0.06, n_wt_ = 9, n_FØ[*skor1a*]_ = 9).

**Figure 3 - figure supplement 3.**
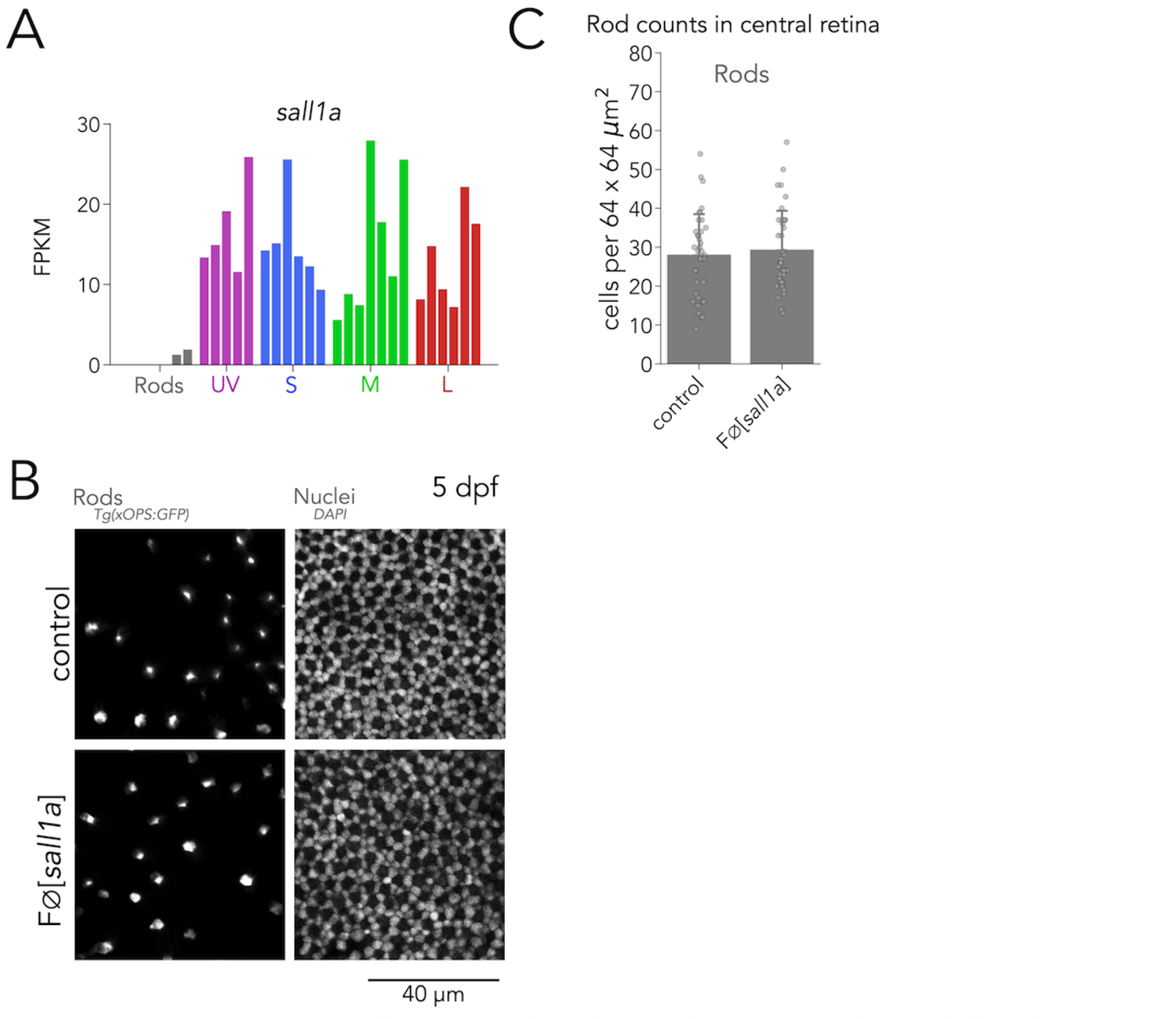
Sall1a is not required for photoreceptor specification. **(A)** Expression of *sall1a* across photoreceptors shows enrichment in all cone subtypes but not rods. **(B)** Mutations in *sall1a* do not change rod densities and the cone mosaic is not disrupted. Representative confocal images of the central retina of control (top row) and FØ[*sall1a*] (bottom row) larvae at 5 dpf. Columns correspond to a transgenic line that rods or to nuclei labeled with DAPI to reveal the cone mosaic. **(C)** Quantification of photoreceptor densities in control and FØ[*sall1a*] larvae. Bars represent averages, error bars correspond to standard deviations, and markers correspond to individual retinas. There is no significant change in the density of rods in FØ[*sall1a*] compared to *wt controls* (Mann-Whitney U = 714.00, p = 0.79, n_wt_ = 38, n_FØ[*sall1a*]_ = 39).

**Figure 3 - figure supplement 4.**
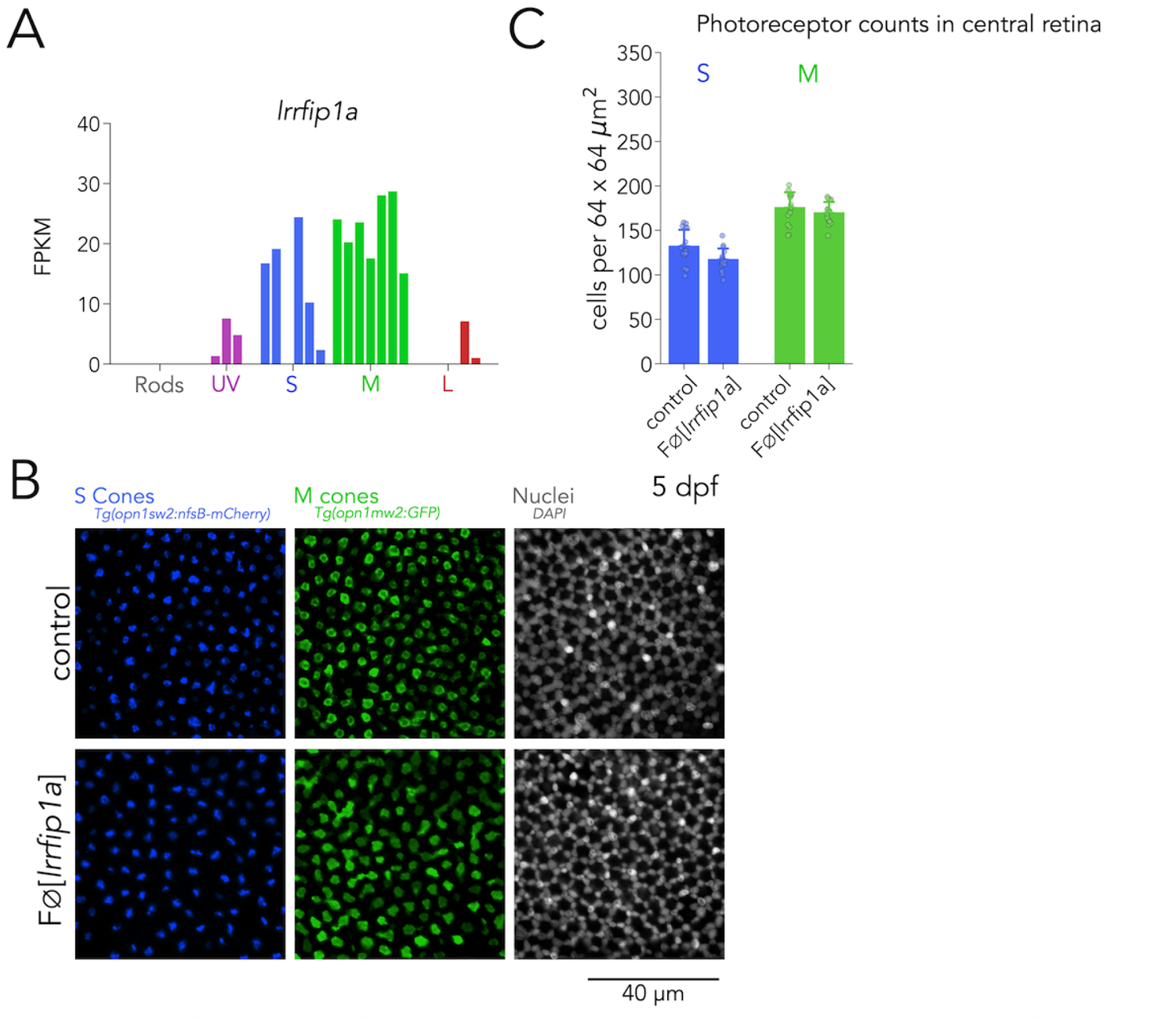
Lrrfip1a is not required for M-cone specification. **(A)** Expression of *lrrfip1a* across photoreceptors shows enrichment in M cones. **(B)** Mutations in *lrrfip1a* do not change S-cone or M-cone densities and the cone mosaic is not disrupted. Representative confocal images of the central retina of control (top row) and FØ[*lrrfip1a*] (bottom row) larvae at 5 dpf. Columns correspond to a transgenic line that S cones (blue) or M cones (green) or to nuclei labeled with DAPI to reveal the cone mosaic. **(C)** Quantification of photoreceptor densities in control and FØ[*lrrfip1a*] larvae. Bars represent averages, error bars correspond to standard deviations, and markers correspond to individual retinas. There is no significant change in the density of M cones in FØ[*lrrfip1a*] compared to wt controls (Mann-Whitney U = 153.50, p = 0.19, n_wt_ = 16, n_FØ[*lrrfip1a*]_ = 15), or in the density of S cones in FØ[*lrrfip1a*] (Mann-Whitney U = 178.00, p = 0.02, n_wt_ = 16, n_FØ[*lrrfip1a*]_ = 15).

**Figure 3 - figure supplement 5.**
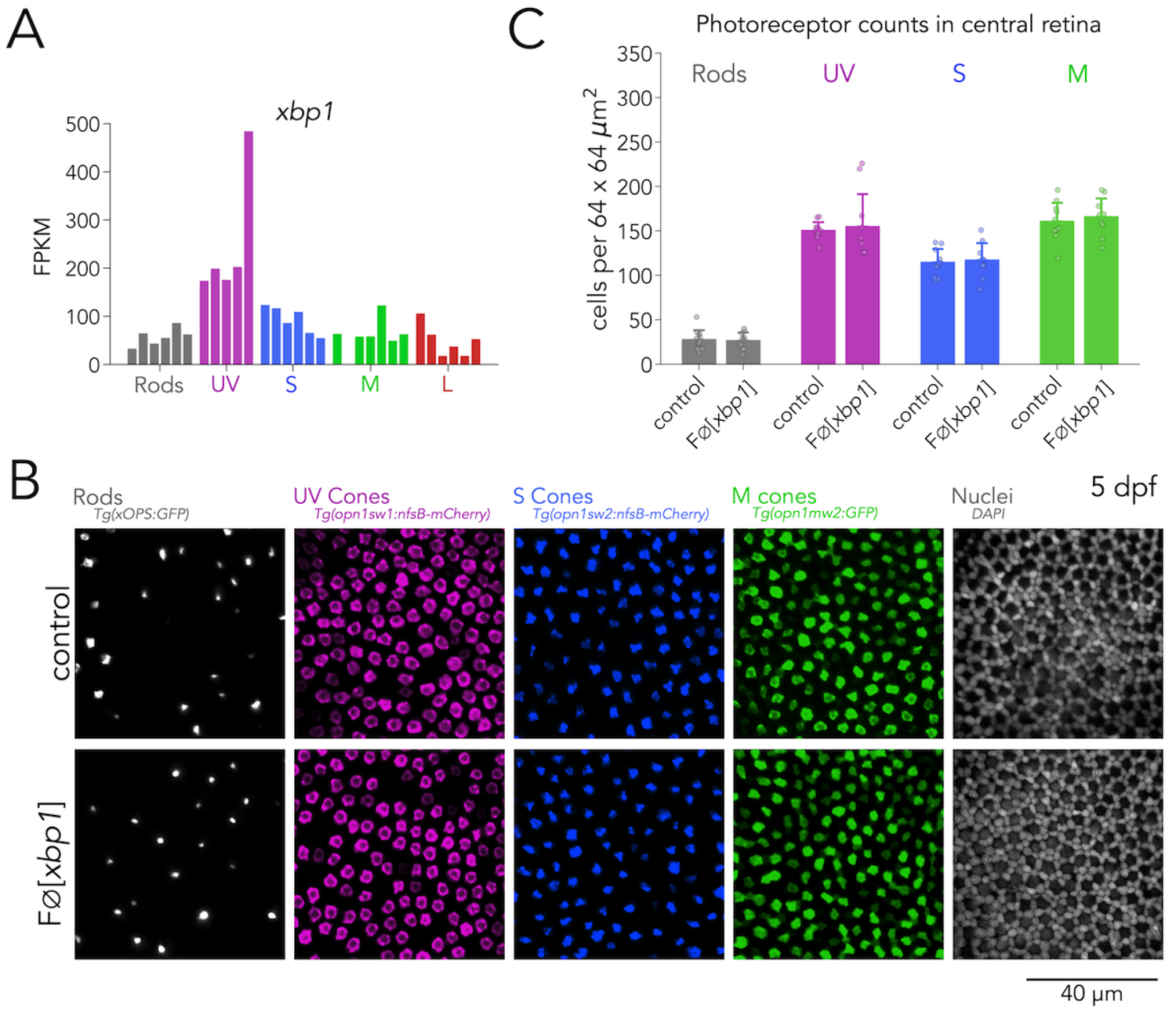
Xbp1 is not required for photoreceptor specification. **(A)** *xbp1* is expressed in all photoreceptor subtypes but with enrichment in UV cones. **(B)** Mutations in *xbp1* do not change photoreceptor densities and cone mosaic is not disrupted. Representative confocal images of the central retina of control (top row) and FØ[*xbp1*] (bottom row) larvae at 5 dpf. Columns correspond to a transgenic line that labels a photoreceptor subtypes or to nuclei labeled with DAPI to reveal the cone mosaic. **(C)** Quantification of photoreceptor densities in control and FØ[*xbp1*] larvae. Bars represent averages, error bars correspond to standard deviations, and markers correspond to individual retinas. There is no significant change in the density of rods in FØ[*xbp1*] (Mann-Whitney U = 54.50, p = 1.00, n_wt_ = 11, n_FØ[*xbp1*]_ = 10), UV cones (Mann-Whitney U = 67.00, p = 0.42, n_wt_ = 11, n_FØ[*skor1a*]_ = 10), S cones (Mann-Whitney U = 40.50, p = 0.74, n_wt_ = 10, n_FØ[*xbp1*]_ = 9), or of M cones (Mann-Whitney U = 39.50, p = 0.68, n_wt_ = 10, n_FØ[*xbp1*]_ = 9) compared to *wt* controls.

**Figure 4 - figure supplement 1.**
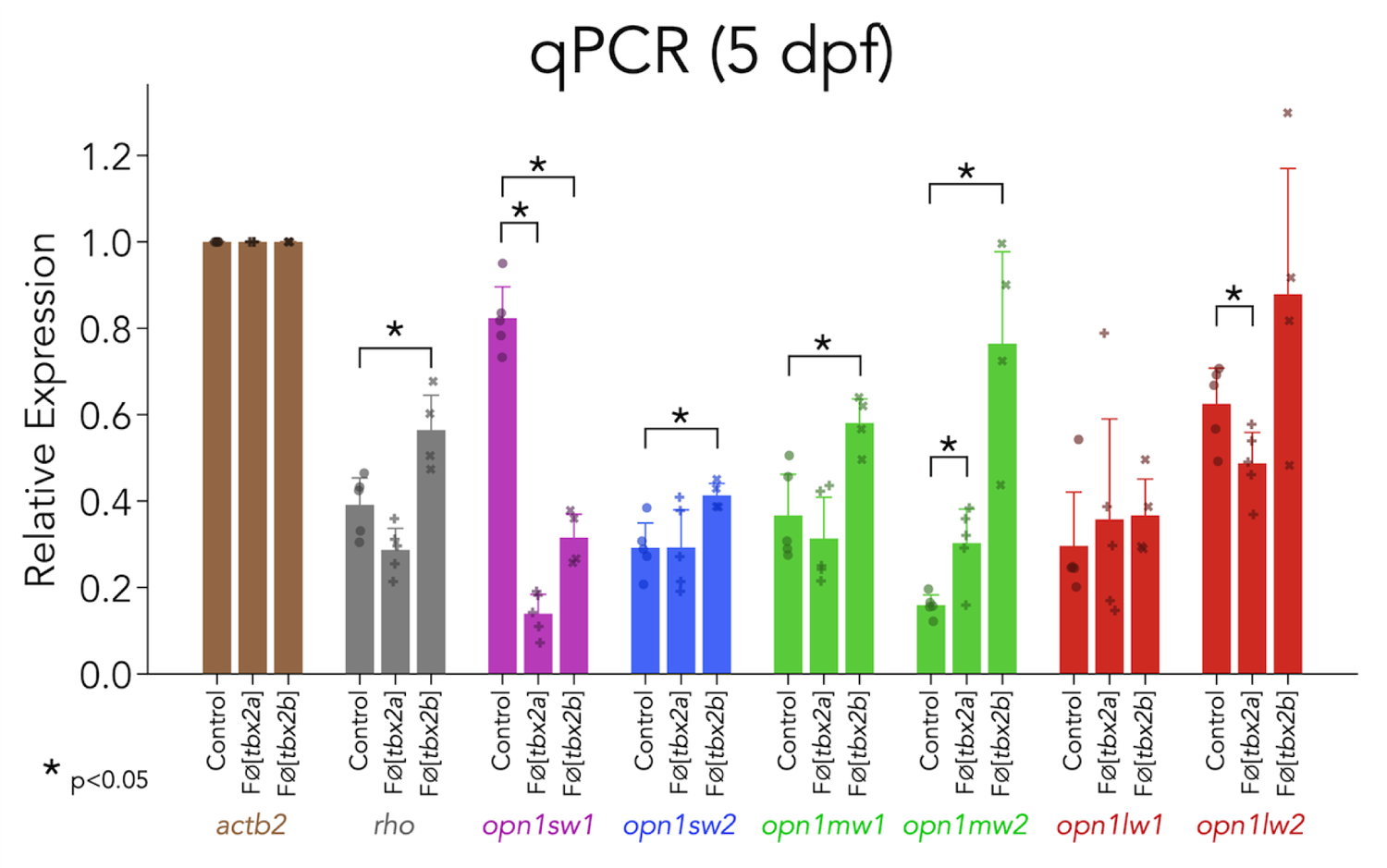
Mutations in *tbx2a* and *tbx2b* cause multiple changes in opsin expression. Quantification of opsin expression using real-time quantitative PCR, using the standard curve method and relative to expression of actin (*actb2*). Compared to controls (n_wt_ = 5), FØ[*tbx2a*] (n_FØ[tbx2a]_ = 5) showed significant changes in the expression of UV opsin (*opn1sw1*, Mann-Whitney U = 25, p = 0.008), M opsin (*opn1mw2*, Mann-Whitney U = 2.0, p = 0.03) and L opsin (*opn1lw2*, Mann-Whitney U = 22, p = 0.05). FØ[*tbx2b*] (n_FØ[tbx2b]_ = 4) showed significant changes in the expression of Rhodopsin (*rho*, Mann-Whitney U = 0, p = 0.01), UV opsin (*opn1sw1*, Mann-Whitney U = 20, p = 0.01), S opsin (*opn1sw2*, Mann-Whitney U = 0, p = 0.01) and M opsin (*opn1mw1*, Mann-Whitney U = 1, p = 0.03; *opn1mw2*, Mann-Whitney U = 0, p = 0.01). We ran all measurements in triplicate. We were not able to detect expression of *opn1mw3* or *opn1mw4*.

